# PCNA ubiquitination protects stalled replication forks from DNA2-mediated degradation by regulating Okazaki fragment maturation and chromatin assembly

**DOI:** 10.1101/759985

**Authors:** Tanay Thakar, Wendy Leung, Claudia M. Nicolae, Kristen E. Clements, Binghui Shen, Anja-Katrin Bielinsky, George-Lucian Moldovan

## Abstract

Upon genotoxic stress, PCNA ubiquitination allows for replication of damaged DNA by recruiting lesion-bypass DNA polymerases. However, PCNA is also ubiquitinated during normal S-phase progression. By employing ubiquitination-deficient 293T and RPE1 cells generated through CRISPR/Cas9 genome editing, we show that this modification promotes cellular proliferation and suppression of genomic instability under normal growth conditions. Loss of PCNA-ubiquitination results in DNA2-mediated but MRE11-independent nucleolytic degradation of nascent DNA at stalled replication forks. This degradation is linked to defective gap-filling in the wake of the replication fork, and incomplete Okazaki fragment synthesis and maturation, thus interfering with efficient PCNA unloading by ATAD5 and subsequent nucleosomal deposition by CAF-1. Moreover, concomitant loss of PCNA-ubiquitination and BRCA2 results in a synergistic increase in nascent DNA degradation and sensitivity to PARP-inhibitors. In conclusion, we show that by ensuring efficient Okazaki fragment maturation, PCNA-ubiquitination protects fork integrity and promotes the resistance of BRCA-deficient cells to PARP-inhibitors.

## Introduction

Accurate DNA replication is essential for genomic stability and suppression of mutagenesis. DNA replication (O’Donnell et al., 2013; Siddiqui et al., 2013) is initiated at discrete replication origins, marked by loading of origin recognition complexes, and proceeds bidirectionally upon back-to-back loading of two copies of the MCM helicase complex. DNA replication occurs in a continuous manner on the leading strand, catalyzed by DNA polymerase Polε, whereas lagging strand replication occurs discontinuously, needing frequent re-priming by the Polα-primase complex, followed by processive DNA synthesis by Polδ. This results in short RNA-primed DNA fragments known as Okazaki fragments (OFs). An essential component of the replication machinery is the homotrimeric ring-shaped protein Proliferating Cell Nuclear Antigen (PCNA), which encircles the DNA molecule, sliding along it as DNA synthesis proceeds. PCNA is loaded at replication origins by the RFC1-5 complex, and unloaded upon replication termination by an alternative complex in which ATAD5 (Elg1 in yeast) replaces RFC1 (Kubota et al., 2013; Lee et al., 2013).

During DNA synthesis, PCNA interacts with the replicative polymerases on each strand and enhances their processivities, rendering PCNA essential for DNA replication and cellular proliferation (Choe and Moldovan, 2017; Leung et al., 2018). On the lagging strand PCNA also recruits and coordinates the activity of the Flap endonuclease (FEN1) which cleaves the RNA primer displaced by Polδ, and DNA ligase 1 (LIG1) which seals the resulting nick to complete OF maturation (OFM) (Zheng and Shen, 2011). Concomitant with DNA replication, PCNA also controls chromatinization of the newly synthesized DNA by recruiting the chromatin assembly factor CAF-1 and other histone chaperone complexes (Alabert et al., 2017; Sauer et al., 2018). In general, these interactions are mediated by a conserved motif termed PCNA-interacting peptide (PIP)-box present on most PCNA binding partners (Choe and Moldovan, 2017; Mailand et al., 2013).

The normal progression of replication forks can be hampered upon encountering unrepaired DNA lesions which block the progression of replicative polymerases, a process known as replication stress (Techer et al., 2017; Zeman and Cimprich, 2014). Genomic regions harboring repetitive elements, DNA secondary structures, and other difficult to replicate sequences, can also induce the arrest of the replicative polymerases, serving as endogenous sources of replication stress. In response to replication stress, PCNA is mono-ubiquitinated by the RAD18 ubiquitin ligase at lysine 164 (K164). This modification promotes a switch from the replicative polymerase to specialized low-fidelity polymerases, which contain both PIP-boxes as well as ubiquitin-binding motifs, and thus preferentially bind to ubiquitinated PCNA (^Ubi^PCNA) (Bienko et al., 2005; Choe and Moldovan, 2017; Guo et al., 2006; Hoege et al., 2002; Kannouche et al., 2004; Leung et al., 2018; Stelter and Ulrich, 2003). These specialized polymerases promote the bypass of replication obstacles in order to maintain efficient DNA-replication, a process known as translesion synthesis (TLS) (Vaisman and Woodgate, 2017; Yang and Gao, 2018).

In response to replication stress, forks can be reversed, which involves their processing into four-way junctions upon annealing of the complementary nascent strands. Fork reversal is thought to function as a protection mechanism against fork collapse by providing an opportunity to bypass the DNA injury by using the nascent strand of the intact sister chromatid as a temporary template for DNA synthesis (Bhat and Cortez, 2018; Cortez, 2019; Quinet et al., 2017). However, reversal can also render replication forks susceptible to nucleolytic processing. In cells lacking a functional BRCA pathway, reversed forks are no longer protected by RAD51 and are subject to excessive resection by the nuclease MRE11 (Schlacher et al., 2011; Schlacher et al., 2012). This nucleolytic degradation drives genome instability and underlies the sensitivity of BRCA-mutant cells to cisplatin and PARP inhibitors (PARPi) (Ray Chaudhuri et al., 2016; Taglialatela et al., 2017). In addition to MRE11, other nucleases, including DNA2, EXO1, CTIP, and MUS81 have been implicated in resection of nascent DNA at stalled forks and subsequent genome instability (Higgs et al., 2015; Lemacon et al., 2017; Rondinelli et al., 2017; Thangavel et al., 2015). Therefore, protection of replication forks from aberrant resection of nascent DNA is crucial for maintaining genome stability.

In vertebrate cells, mono-ubiquitination is the prevalent form of modified PCNA, although poly-ubiquitination can also be detected (Arakawa et al., 2006; Motegi et al., 2008; Unk et al., 2008). While PCNA ubiquitination is induced upon replication stress, basal levels of mono-ubiquitinated PCNA can be detected in S-phase cells under unperturbed growth conditions (Arakawa et al., 2006; Kannouche et al., 2004; Unk et al., 2008). This suggests that, in human cells, PCNA ubiquitination may play an important but so far elusive role in controlling replication fork progression and genome stability during normal S-phase. Here, we show that loss of PCNA ubiquitination renders nascent DNA at stalled replication forks susceptible to degradation by the nuclease DNA2. Mechanistically, we link this nucleolytic degradation to defective gap-filling in the wake of the replication fork and demonstrate that the inability to ubiquitinate PCNA at K164 results in the same phenotypes as aberrant OFM. Defective OFM and the PCNA-K164R mutation are epistatic and both preclude efficient PCNA unloading by ATAD5, which subsequently impairs nucleosomal deposition by CAF-1. Moreover, we demonstrate that loss of PCNA ubiquitination results in a synergistic increase in nascent strand degradation and genomic instability in BRCA2-deficient cells, and enhances PARPi sensitivity of these cells. We therefore define the ^Ubi^PCNA–LIG1–ATAD5–CAF-1 pathway as a novel genetic axis protecting replication fork stability and maintaining the integrity of the genome that operates in parallel to the BRCA-RAD51-MRE11 pathway.

## Results

### Generation of PCNA-K164R mutant cells

As PCNA is essential for cell proliferation, previous studies investigating the role of PCNA ubiquitination in human cell lines heavily relied on siRNA-mediated depletion of endogenous PCNA coupled with transfection of a K164R mutant or PCNA-ubiquitin fusion polypeptides (Leung et al., 2018). However, the residual expression from the endogenous PCNA locus and the artificial overexpression of the PCNA variants can complicate the analyses. In order to overcome these limitations, we employed the CRISPR/Cas9 genome editing system to introduce the K164R homozygous mutation in the endogenous PCNA gene, in 293T and RPE1 cell lines. Monoclonal cultures were initially screened for loss of PCNA ubiquitination by western blot using an antibody specific for ubiquitinated PCNA. Several ubiquitination-deficient 293T clones were obtained. However, we noticed that in these clones, the level of unmodified PCNA was reduced compared to the parental line, as shown for one of the clones, designated KR5, in Figure S1A. The genome of 293T cells is considered pseudotriploid (Lin et al., 2014). Upon cloning of individual PCNA alleles from the KR5 cell line, followed by Sanger sequencing, we found that a single PCNA allele was edited with the desired mutation, while the others were inactivated through introduction of small insertions or deletions. To exclude phenotypes caused by reduced PCNA expression and other potential off-target effects of the CRISPR/Cas9 procedure, we created an isogenic pair by re-expressing either wildtype or a K164R mutant of PCNA in the KR5 clone through a lentiviral expression system. The resulting cell lines, termed 293T-WT and 293T-K164R (or KR) from here on, show similar levels of unmodified PCNA between themselves and when compared to the parental cell line (Figure 1A; Figure S1A), and were thus used for subsequent studies.

**Figure 1.**
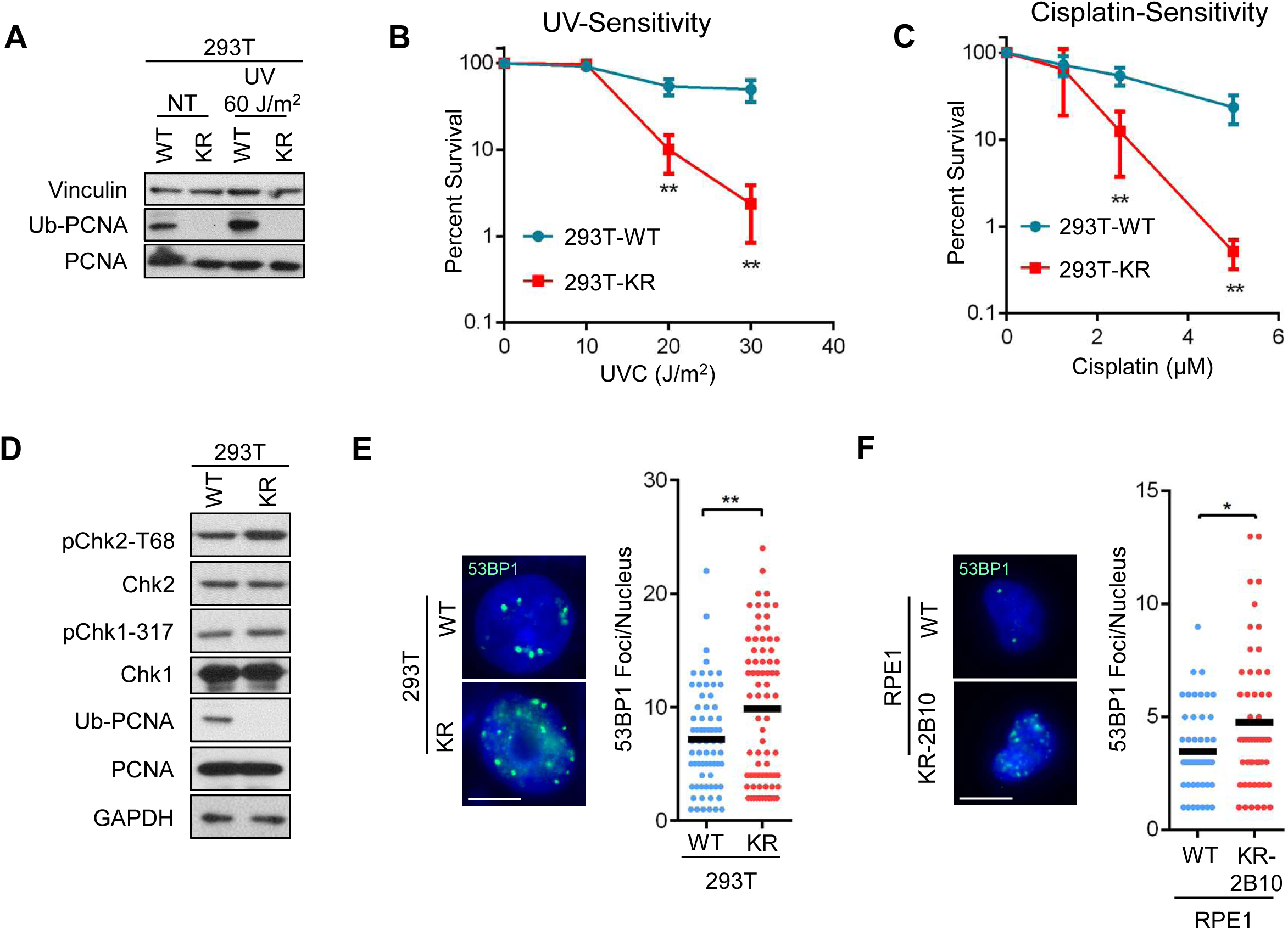
PCNA ubiquitination-deficiency leads to accumulation of DNA damage under normal growth conditions. **A.** Western blot experiment showing the loss of PCNA ubiquitination in 293T-K164R cells generated through CRISPR/Cas9 genome editing. Denatured whole cell extracts of cells under normal growth conditions, or 3h after exposure to the indicated UV dose, were analyzed. A similar experiment performed in RPE1-K164R cells is shown in Figure S1. **B, C**. Clonogenic survival experiments showing hypersensitivity of 293T-K164R cells to UV (**B**) and cisplatin (**C**). The average of three experiments for UV and four experiments for cisplatin, with standard deviations indicated as error bars, is shown. Asterisks indicate statistical significance. **D**. Western blot experiment showing increased Chk2 phosphorylation in 293T-K164R cells under normal growth conditions. **E, F**. Immunofluorescence experiment showing increased 53BP1 chromatin foci in unsynchronized 293T-K164R (**E**) and RPE1-K164R (**F**) cells. At least 50 cells were quantified for each condition. The mean value is represented on the graphs, and asterisks indicate statistical significance. Representative micrographs are also shown.

In contrast to 293T cells, RPE1 cells are nearly diploid (Mardin et al., 2015). Two RPE1 clones expressing an endogenous PCNA-K164R mutant were generated (Figure S1B). Both mutant clones showed similar levels of unmodified PCNA as the parental line. Sequencing of the genomic region targeted confirmed that, in both clones, both PCNA alleles were homozygously edited with the desired mutation, and thus they were used as such for subsequent experiments (without the complementation employed for 293T cells as described above).

### Endogenous replication stress and increased fork speed in PCNA-K164R cells

As expected from the well-established role of PCNA in TLS, 293T-K164R cells were sensitive to DNA damaging agents that induce single-stranded DNA lesions, such as UV and cisplatin (Figure 1B, C). Moreover, 293T-K164R cells showed reduced UV-induced mutagenesis rates (Figure S1C) as measured by the SupF shuttle vector assay (Wang et al., 1995) –in line with the role of PCNA ubiquitination in recruiting the TLS polymerase Polη to bypass UV-induced lesions (Kannouche et al., 2004). Rather unexpectedly, however, under unperturbed growth conditions, KR clones showed lower proliferation rates (Figure S1D), coupled with a reduced proportion of cells undergoing DNA synthesis as measured by EdU incorporation (Figure S1E). As this pattern was reminiscent of cells experiencing increased levels of endogenous replication stress (Daigh et al., 2018), we next investigated expression of DNA damage markers. We observed that, in the absence of any exogenous DNA damage treatment, KR cells showed increased levels of Chk2 phosphorylation (Figure 1D) and 53BP1 chromatin foci (Figure 1E-F), indicating DSB accumulation. These findings suggest that PCNA ubiquitination-deficient cells are unable to resolve endogenous replication stress, resulting in DNA damage accumulation under unperturbed growth conditions.

To evaluate the role of PCNA ubiquitination in replication fork progression, we employed the DNA fiber combing assay to measure replication dynamics in 293T-K164R and RPE1-K164R cells. In this assay, nascent replication tracts can be identified and quantified upon consecutive incubation of cells with the thymidine analogs IdU and CldU, followed by combing of intact DNA molecules on glass coverslips and immunofluorescence-based detection of the incorporated nucleoside analogs with specific antibodies. Under unperturbed growth conditions, 293T-K164R cells showed longer nascent tract length and increased replication fork speed (Figure 2A; Figure S2A). Previously, an inverse correlation between origin firing and fork speed was reported (Maya-Mendoza et al., 2018; Zhong et al., 2013). In line with this, we observed reduced origin firing in 293T-K164R cells (Figure S2B). Similar results were obtained in RPE1-K164R cells (Figure 2B; Figure S2C, D). The ZRANB3 translocase was previously shown to be recruited by K63-linked polyubiquitinated PCNA to mediate slowing of replication forks in the presence of replication stress (Vujanovic et al., 2017). Indeed, 293T-K164R cells were unable to efficiently reduce fork speed in the presence of low levels (0.4mM) of the replication fork stalling agent hydroxyurea (HU) (Figure S2E). This raises the possibility that the longer nascent tracts observed in KR cells under normal growth conditions may simply reflect the loss of ZRANB3 recruitment to stressed replication forks. To address this, we depleted ZRANB3. Loss of the protein in wildtype cells did not result in longer nascent tracts (Figure S2F, G), arguing against a role for ZRANB3-mediated fork slowing in controlling fork speed under normal growth conditions.

**Figure 2.**
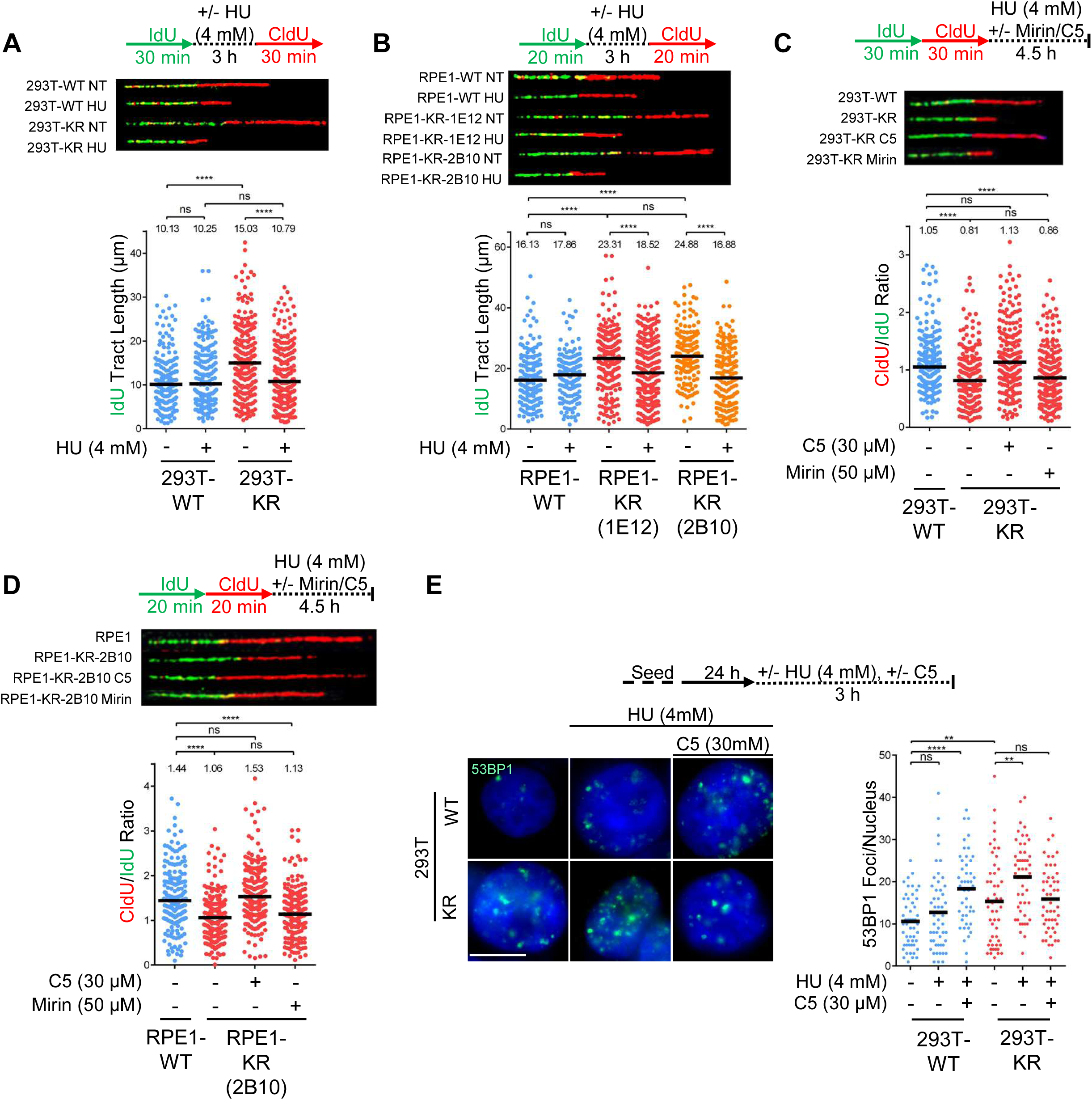
PCNA-K164R cells exhibit DNA2-mediated, but MRE11-independent nascent DNA degradation upon replication arrest. **A, B.** DNA fiber combing assays showing faster replication fork progression in 293T-K164R cells (**A**) and two different clones of RPE1-K164R cells (**B**) under normal growth conditions, and nascent strand degradation upon HU treatment. The quantification of the IdU tract length is presented, with the median values marked on the graph and listed at the top. Asterisks indicate statistical significance. Schematic representation of the assay conditions, and representative micrographs are also presented. **C, D.** HU-induced nascent strand degradation in 293T-K164R (**C**) and RPE1-K164R (**D**) cells is suppressed by incubation with the DNA2 inhibitor C5, but not by treatment with the MRE11 inhibitor mirin. The ratio of CldU to IdU tract lengths is presented, with the median values marked on the graph and listed at the top. Asterisks indicate statistical significance. A schematic representation of the assay conditions, as well as representative micrographs are also presented. **E**. Immunofluorescence experiment showing that HU treatment augments 53BP1 foci formation in unsynchronized 293T-K164R cells. DNA2 inhibition suppresses 53BP1 foci formation in KR, but not in wildtype cells. At least 40 cells were quantified for each condition. The mean value is represented on the graphs, and asterisks indicate statistical significance. Representative micrographs are also shown.

### PCNA ubiquitination protects stalled replication forks against DNA2-mediated nucleolytic degradation

To investigate if the abnormal replication fork characteristics described above are associated with defects in fork stability, we measured replication fork integrity in the presence of acute replication stress. Treatment with 4mM HU resulted in degradation of the nascent DNA tract in both 293T-K164R and RPE1-K164R cells, but not in the respective control cells (Figure 2A-D). Nascent tract degradation was observed under two different experimental conditions: when HU was added for 3h in between the IdU and CldU pulses and the IdU tract length was measured (Figure 2A, B), and when HU was added for 4.5h after consecutive incubations with thymidine analogs and the ratio of CldU to IdU tract-lengths was calculated (Figure 2C, D). HU-induced nascent strand degradation has been extensively described in the context of BRCA deficiency, where it is dependent on the activity of the MRE11 nuclease (Bhat and Cortez, 2018; Schlacher et al., 2011; Schlacher et al., 2012). Surprisingly, MRE11 inhibition using the inhibitor mirin did not suppress nascent tract degradation in KR cells (Figure 2C, D, Figure S2H-J), indicating that a different fork degradation pathway operates in these cells. We further ruled out the involvement of other nucleases previously involved in nascent tract degradation, including EXO1, CTIP, and MUS81 (Lemacon et al., 2017; Rondinelli et al., 2017) (Figure S2K-N). In contrast, inhibition of the nuclease DNA2 with the specific inhibitor C5 completely restored nascent tract length in both 293T-K164R and RPE1-K164R cells (Figure 2C, D; Figure S2J, K), indicating that DNA2 is the nuclease responsible for fork degradation upon loss of PCNA ubiquitination. WRN helicase has been previously described as a cofactor for DNA2 in nascent tract degradation (Thangavel et al., 2015). In line with this, WRN depletion also rescued HU-induced fork degradation in KR cells (Figure S2O, P).

The K164 residue of PCNA is subjected not only to ubiquitination, but also to SUMOylation (Li et al., 2018; Moldovan et al., 2012). Depletion of the ubiquitin ligase RAD18 recapitulated the PCNA-K164R phenotype, as it resulted in DNA2-mediated nascent tract degradation. Importantly, depletion of the SUMO-conjugating enzyme UBC9 did not affect fork stability (Figure S2Q-S), indicating that K164 modification by ubiquitin, rather than SUMO, is necessary for replication fork protection.

Next, we investigated the impact of DNA2-mediated fork degradation on genomic instability in KR cells. HU treatment induced 53BP1 foci preferentially in KR cells compared to WT. DNA2 inhibition suppressed HU-induced 53BP1 foci formation in KR, but not in WT cells (Figure 2E). Similar results were obtained for RPA foci (Figure S2T). These findings argue that, in KR cells, DNA2-mediated processing of stalled replication forks results in DSB formation and genomic instability.

### Nascent tract degradation in PCNA ubiquitination-defective cells depends on fork reversal but is not caused by a defect in fork protection by RAD51

In BRCA-deficient cells, nascent tract degradation by MRE11 occurs upon reversal of stalled replication forks (Mijic et al., 2017; Ray Chaudhuri et al., 2016; Taglialatela et al., 2017). Thus, we investigated if fork reversal is also required for nascent tract degradation in KR cells. Fork reversal depends on RAD51 and the translocases HLTF, ZRANB3, and SMARCAL1 (Cortez, 2019; Kolinjivadi et al., 2017; Mijic et al., 2017; Quinet et al., 2017; Taglialatela et al., 2017; Zellweger et al., 2015). Depletion of RAD51 restored nascent tract integrity in KR cells (Figure 3A; Figure S3A), indicating that fork reversal by RAD51 is indeed a prerequisite for nascent strand degradation in KR cells as well. We next investigated the involvement of translocases HLTF, ZRANB3 and SMARCAL1. Previously, individual depletion of each of these factors was shown to completely restore fork protection in BRCA-deficient cells, suggesting that they act in concert to perform fork reversal (Poole and Cortez, 2017; Taglialatela et al., 2017). In contrast, in KR cells we observed differential impact of translocases depletion. Loss of HLTF did not restore fork protection in KR cells, whereas ZRANB3 depletion partially rescued nascent tract degradation, and complete rescue was only observed after depleting SMARCAL1 (Figure 3B; Figure S3B). These findings demonstrate that, unlike in BRCA-mutant cells, fork reversal in KR cells depends on SMARCAL1 and partially on ZRANB3, and does not involve HLTF activity.

**Figure 3.**
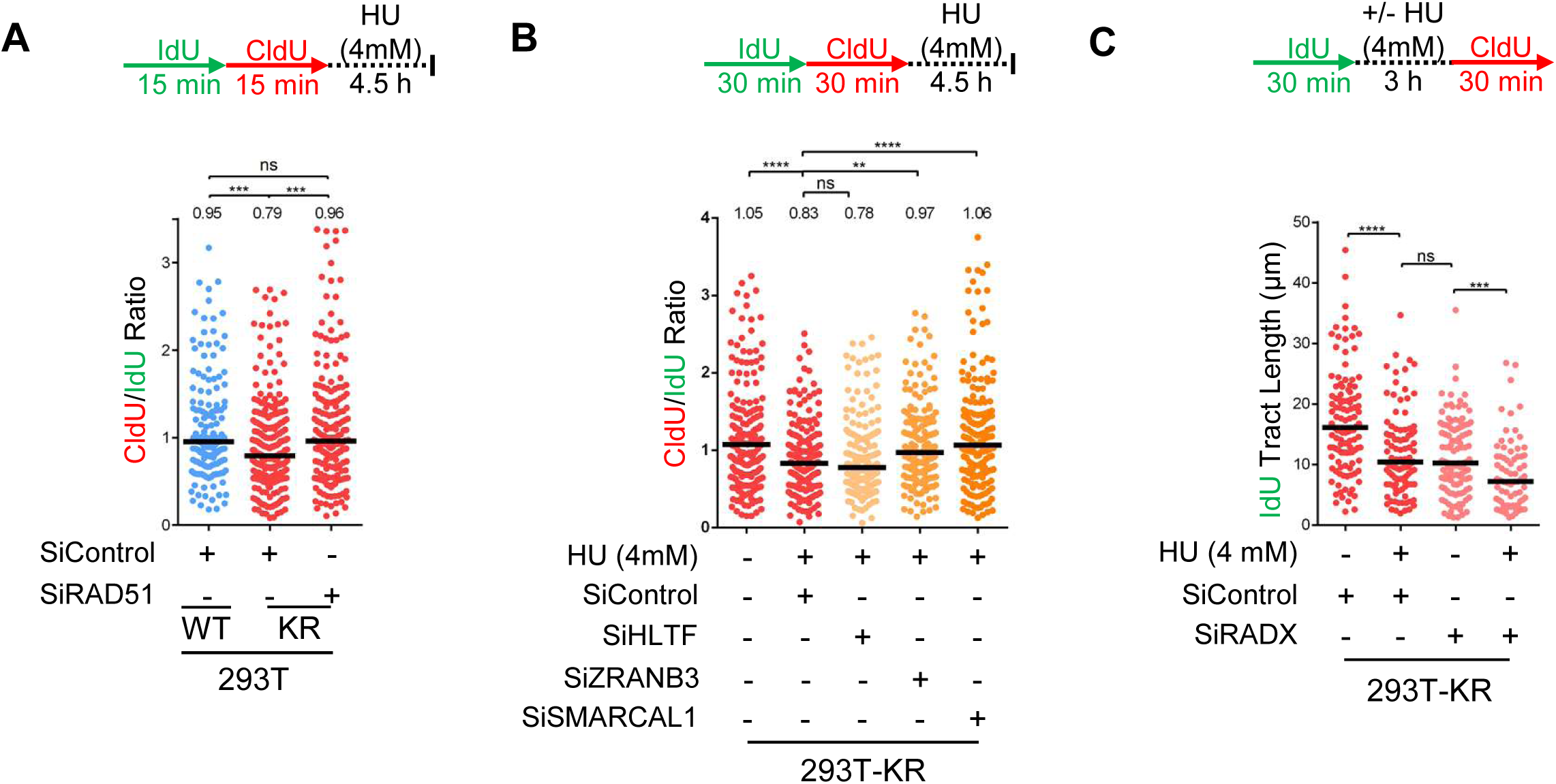
Reversal of stalled forks is necessary for nascent strand degradation in PCNA-K164R mutant cells. **A.** RAD51 depletion suppresses HU-induced nascent strand degradation in 293T-K164R cells. The ratio of CldU to IdU tract lengths is presented, with the median values marked on the graph and listed at the top. Asterisks indicate statistical significance. A schematic representation of the fiber combing assay conditions is also presented. A Western blot showing RAD51 levels upon siRNA treatment is presented in Figure S3A. **B.** Impact of DNA translocases HLTF, ZRANB3, and SMARCAL1 on HU-induced nascent strand degradation in 293T-K164R cells. The ratio of CldU to IdU tract lengths is presented, with the median values marked on the graph and listed at the top. Asterisks indicate statistical significance. A schematic representation of the DNA fiber combing assay conditions is also presented. Western blots confirming the knockdowns are shown in Figure S3B. **C.** Knockdown of RADX does not suppress HU-induced nascent strand degradation in 293T-K164R cells. The quantification of the IdU tract length is presented, with the median values marked on the graph. While RADX depletion results in a reduction in fork progression rate, HU treatment results in further reduction in nascent tract length in both wildtype and KR cells. Asterisks indicate statistical significance. A schematic representation of the assay conditions is also presented. Confirmation of RADX knockdown is shown in Figure S3C.

Besides its role in fork reversal, RAD51 is also critical for the protection of reversed forks. The inability to stabilize RAD51 at stalled replication forks renders them susceptible to nucleolytic processing (Higgs et al., 2015; Schlacher et al., 2011; Schlacher et al., 2012). To test if the fork protection defect observed in KR cells is caused by defective RAD51 loading on reversed forks, we depleted RADX. RADX antagonizes RAD51 accumulation at stalled forks, and its depletion results in enhanced RAD51 binding to reversed forks (Dungrawala et al., 2017). However, RADX knockdown failed to restore fork protection in KR cells (Figure 3C; Figure S3C), arguing that the nascent tract degradation in KR cells is not caused by deficient RAD51-mediated fork protection.

Previously, DNA2-mediated nascent tract degradation was described in cells depleted of the helicase RECQL1, which restarts stalled replication forks, upon prolonged fork arrest (treatment with 4mM HU for 6h) (Thangavel et al., 2015). To test if the fork protection defect in KR cells is caused by defective RECQL1-mediated fork restart, we depleted RECQL1 in both WT and KR cells. Under experimental conditions used to detect nascent tract degradation in KR cells (4mM HU for 3h), we did not observe any impact of RECQL1 knockdown in either WT or KR cells (Figure S3D, E). These findings argue against an involvement of RECQL1 in the fork protection defect observed in KR cells.

### Okazaki fragment ligation and PCNA ubiquitination operate in the same fork protection pathway

The ∼30% increase in fork speed observed in KR cells is reminiscent of cells depleted of DNA replication factors involved in Okazaki fragment maturation (OFM) such as FEN1 and LIG1 (Maya-Mendoza et al., 2018). Thus, we next investigated if perturbing OFM also results in nascent DNA degradation upon fork stalling. Under identical conditions (4mM HU for 3h), LIG1 depletion also induced nascent tract degradation (Figure 4A; Figure S4A). Recently, the PARP1-XRCC1-LIG3 pathway was described as a novel OFM mechanism that operates in parallel to the FEN1-LIG1 pathway (Hanzlikova et al., 2018). PARP1 depletion also resulted in faster replication forks under normal conditions, which were nucleolytically degraded upon acute replication stress (Figure 4B; Figure S4B). These findings indicate that defects in OF maturation result in nascent DNA degradation upon replication stress.

**Figure 4.**
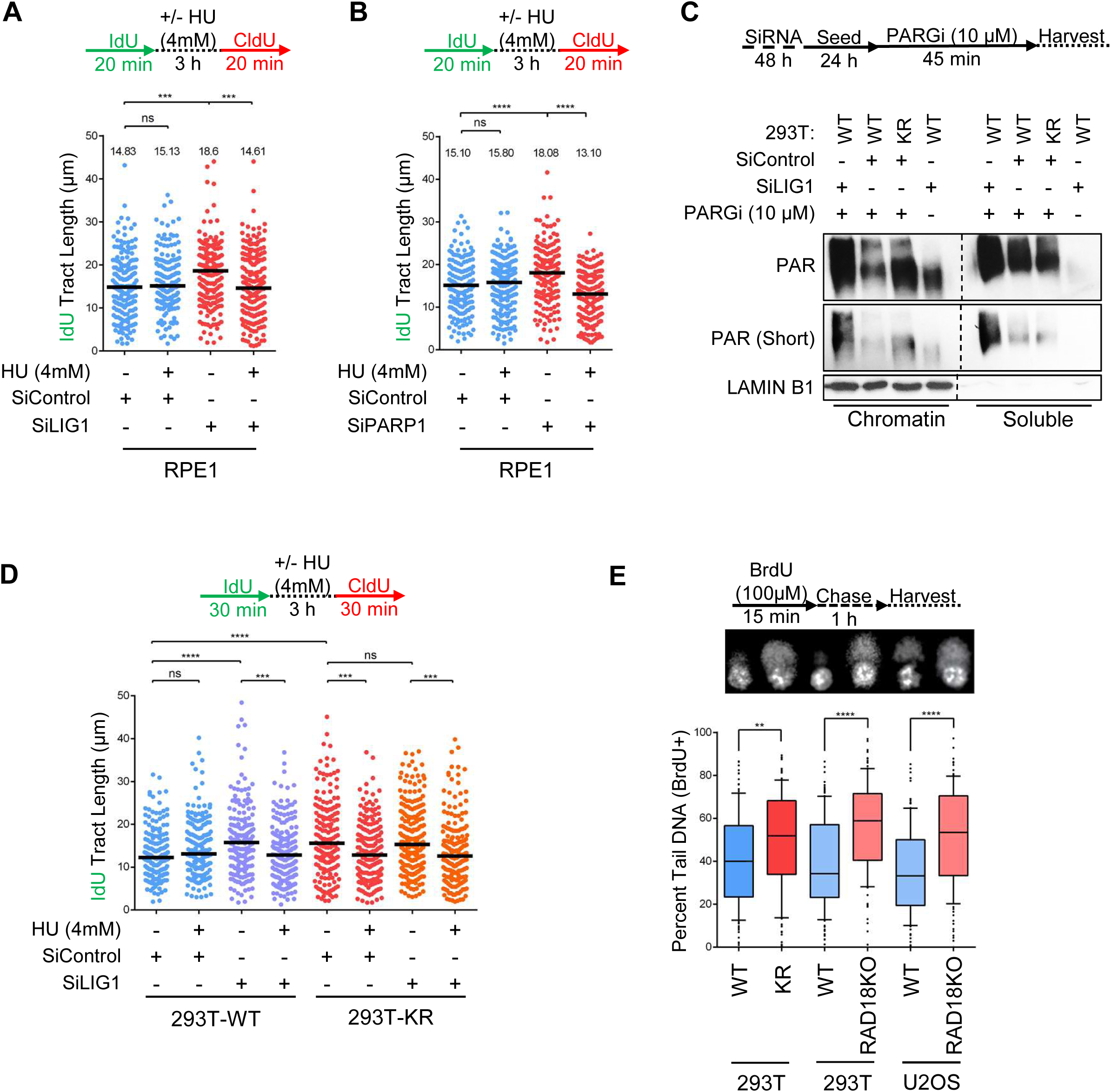
Defective Okazaki fragment maturation results in nascent strand degradation upon replication stress. **A, B.** DNA fiber combing assay showing that LIG1 (**A**) or PARP1 (**B**) depletions result in faster replication fork progression under normal conditions, and induce nascent strand degradation upon fork arrest. The quantification of the IdU tract length is presented, with the median values marked on the graph and listed at the top. Asterisks indicate statistical significance. A schematic representation of the assay conditions is also presented. Western blots confirming LIG1 and PARP1 knockdowns are shown in Figure S4A and Figure S4B, respectively. **C**. Chromatin fractionation experiment showing increased PAR chain formation in KR cells under normal growth conditions, indicating defective Okazaki fragment maturation. Cells were treated as indicated with a PARG inhibitor (PARGi) for 45min prior to harvesting to block PAR chain removal. LIG1 depletion was used as positive control for defective Okazaki fragment maturation. Chromatin-associated laminB1 was used as loading control. **D.** Loss of LIG1 is epistatic with the PCNA-K164R mutation for fork progression and HU-induced nascent strand degradation in 293T cells. The quantification of the IdU tract length is presented, with the median values marked on the graph. While LIG1 knockdown in wildtype cells results in faster fork progression under normal conditions, and HU-induced nascent strand degradation, it does not further increase these phenotypes in KR cells. Asterisks indicate statistical significance. A schematic representation of the assay conditions is also presented. Similar results in RPE1 cells are presented in Figure S4C. **E**. BrdU-alkaline comet assay showing accumulation of ssDNA gaps under normal replication conditions in 293T-K164R and RAD18-knockout 293T and U2OS cells. At least 100 cells were quantified for each condition. The median values are indicated. Asterisks indicate statistical significance comparing means. A schematic representation of the assay conditions, and representative micrographs, are also presented. Western blots confirming RAD18 knockout are shown in Figure S4D.

Using synthetic genetic array analyses in yeast, we previously uncovered a striking genetic similarity between the PCNA-K164R mutation and inactivation of lagging strand synthesis factors (Becker et al., 2015). Coupled with the fork protection defect similarities described above, these findings raise the question of whether PCNA-K164R cells have defects in OF maturation. Previously, it was shown that LIG1 depletion in human cells results in increased poly-ADP-ribose (PAR) chain formation on chromatin (Hanzlikova et al., 2018). We found that PAR chain formation is also enhanced in KR cells (Figure 4C), indicating that PCNA ubiquitination is required for efficient LIG1-mediated Okazaki fragment ligation. Moreover, in both 293T and RPE1 cells, depletion of LIG1 induced fork degradation in WT cells as described above, but did not further exacerbate nascent strand degradation, nor did it further increase fork speed, in KR cells (Figure 4D; Figure S4C). This epistasis indicates that LIG1-mediated OFM, and PCNA ubiquitination may operate in the same fork protection pathway.

Because of the well described roles of PCNA ubiquitination in recruitment of non-canonical polymerases to DNA, we hypothesized that upon endogenous replication stress, the inability to recruit specialized polymerases in KR cells may result in accumulation of single stranded gaps which could hinder OF ligation on the lagging strand. To test this, we performed an alkaline comet assay on cells labeled with BrdU, allowing us to specifically detect single stranded gaps in newly replicated DNA. KR cells showed an increase in DNA gap formation under normal replication conditions, as did cells harboring CRISPR/Cas9-mediated knockout of RAD18 (Figure 4E; Figure S4D). Previously, the TLS polymerase Polκ was shown to be recruited by ubiquitinated PCNA to perform gap filling during nucleotide excision repair (NER) (Ogi et al., 2010). Thus, we tested if Polκ is involved in fork protection similarly to PCNA ubiquitination. POLK depletion resulted in nascent tract degradation, which was partially dependent on DNA2 (Figure S4E, F). In contrast, depletion of the REV1 polymerase did not affect fork stability (Figure S4G, H). These findings show that PCNA ubiquitination suppresses accumulation of under-replicated DNA, which may otherwise interfere with OF ligation on the lagging strand.

### Abnormal PCNA retention on chromatin drives nascent tract degradation through altered nucleosome deposition

We next investigated how OFM defects interfere with fork stability. Previous work in yeast showed that PCNA is removed from DNA by the unloader Elg1 upon DNA Ligase I (Cdc9 in yeast) –mediated sealing of OFs (Kubota et al., 2015). Indeed, knockdown of the Elg1 human homolog ATAD5 or of LIG1 in 293T cells resulted in increased number of PCNA chromatin foci (Figure 5A, Figure S5A, B). Importantly, 293T-K164R cells also showed increased number of PCNA chromatin foci under otherwise unperturbed conditions (Figure 5A), indicating prolonged PCNA retention on chromatin. We next tested if this retention may be responsible for the fork protection defect observed. DNA fiber combing experiments showed that, similar to LIG1 depletion, ATAD5 knockdown in WT cells results in HU-induced degradation of the nascent strand by DNA2 (Figure 5B). Moreover, ATAD5 depletion in KR cells does not further exacerbate the fork protection defect, indicating that ATAD5 participates in the ^Ubi^PCNA–LIG1 pathway of fork protection. These findings show that increased retention of PCNA on chromatin caused by defects in OF ligation or PCNA unloading, results in nascent tract degradation upon fork arrest and reversal.

**Figure 5.**
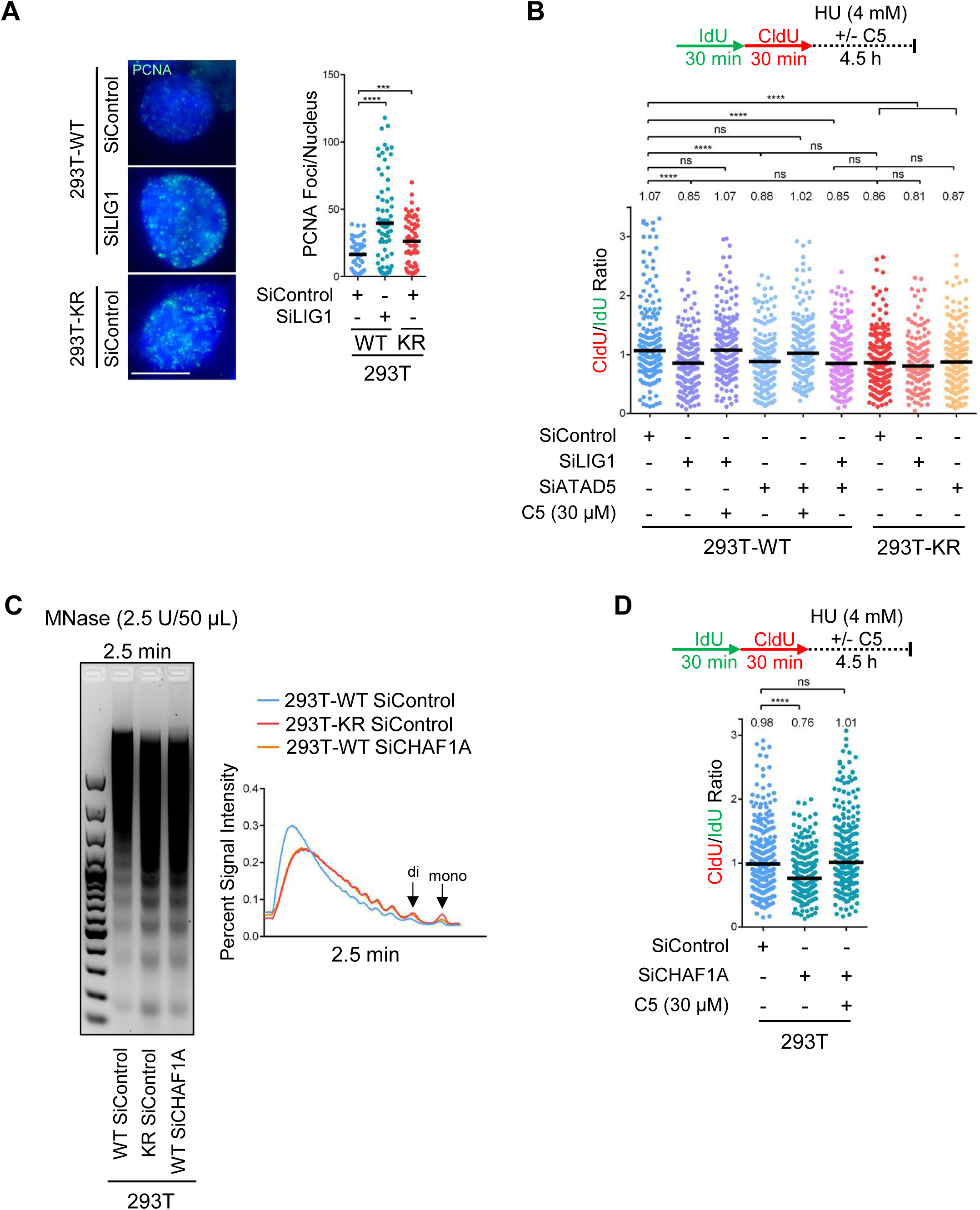
Abnormal retention of PCNA on chromatin upon defective Okazaki fragment maturation drives replication fork degradation by altering nucleosome deposition. **A.** PCNA immunofluorescence showing increased PCNA retention on chromatin in 293T-K164R cells, or upon LIG1 depletion. At least 65 cells were quantified for each condition. The mean value is represented on the graphs, and asterisks indicate statistical significance. Representative micrographs are also shown. **B**. Similar to LIG1 depletion, ATAD5 knockdown results in DNA2-mediated nascent strand degradation upon HU-induced replication fork arrest, which is epistatic to the K164R mutation. The ratio of CldU to IdU tract lengths is presented, with the median values marked on the graph and listed at the top. ATAD5 and LIG1 depletion in wildtype cells results in fork degradation, while their depletion in K164R cells does not further increase the degradation observed in these cells. Asterisks indicate statistical significance. A schematic representation of the fiber combing assay conditions is also presented. Confirmation of ATAD5 knockdown is shown in Figure S5B. **C**. Microccocal nuclease sensitivity assay showing altered nucleosomal deposition in 293-K164R cells, similar to that observed upon depletion of the chromatin assembly factor CHAF1A. A quantification of the signal intensity is also shown. Confirmation of CHAF1A knockdown is shown in Figure S5C. **D**. CHAF1A depletion results in HU-induced degradation of nascent DNA by DNA2 nuclease. The ratio of CldU to IdU tract lengths is presented, with the median values marked on the graph and listed at the top. Asterisks indicate statistical significance. A schematic representation of the fiber combing assay conditions is also presented.

Elg1-deficient yeast cells exhibit nucleosomal assembly defects, as PCNA retention on chromatin also sequesters its binding partner, the histone chaperone complex CAF-1 involved in co-replicational nucleosome assembly (Janke et al., 2018). Thus, we next investigated if nucleosome deposition is altered upon inactivation of the ^Ubi^PCNA–LIG1– ATAD5 pathway. Micrococcal nuclease (MNase) sensitivity assays showed that chromatin was more accessible to nucleolytic digestion in KR cells compared to control cells (Figure 5C), consistent with a reduction in chromatin compaction (Schwab et al., 2013). This defective nucleosome packaging mirrors that observed upon depletion of the CAF-1 complex subunit CHAF1A (Figure 5C; Figure S5C), suggesting that retention of CAF-1 by PCNA in KR cells sequesters CAF-1 away from active nucleosome deposition sites. We therefore investigated the impact of CAF-1 on fork stability. CHAF1A depletion resulted in DNA2-mediated nascent tract degradation upon HU treatment (Figure 5D), similar to what we previously observed upon inactivation of the ^Ubi^PCNA–LIG1–ATAD5 pathway. Altogether, these findings indicate that in KR cells, enhanced PCNA retention on chromatin results in altered nucleosomal packaging likely due to aberrant CAF-1 localization. This nucleosomal packaging defect renders stalled forks susceptible to DNA2-mediated degradation under acute replication stress.

### PCNA ubiquitination and the BRCA pathway act synergistically to protect stalled forks and ensure genomic stability and PARPi resistance

In BRCA-deficient cells, fork protection correlates with resistance to cisplatin and PARPi (Dungrawala et al., 2017; Ray Chaudhuri et al., 2016; Taglialatela et al., 2017). As our findings indicate that PCNA ubiquitination controls a parallel pathway of fork stability to the BRCA pathway, we hypothesized that BRCA inactivation in PCNA-K164R cells will dramatically increase nascent DNA degradation. Indeed, in contrast to LIG1 knockdown, BRCA2 depletion in KR cells resulted in a synergistic increase in nascent tract degradation upon HU treatment (Figure 6A), indicating that the two pathways are operating separately to maintain fork stability. Moreover, a synergistic increase in 53BP1 foci was observed in BRCA2-depleted KR cells under normal growth conditions (Figure 6B), indicating that this fork stability defect translates into DNA damage accumulation and genomic instability. Finally, we investigated the impact of PCNA ubiquitination on the response to PARPi, to which BRCA-deficient cells are hypersensitive (Bryant et al., 2005; Farmer et al., 2005). While the KR mutation by itself did not result in olaparib sensitivity, it enhanced the sensitivity of BRCA1- or BRCA2-depleted cells (Figure 6C-E). These findings show that PCNA ubiquitination provides an alternative mechanism for fork protection in BRCA-deficient cells, and contributes to PARPi resistance in these cells.

**Figure 6.**
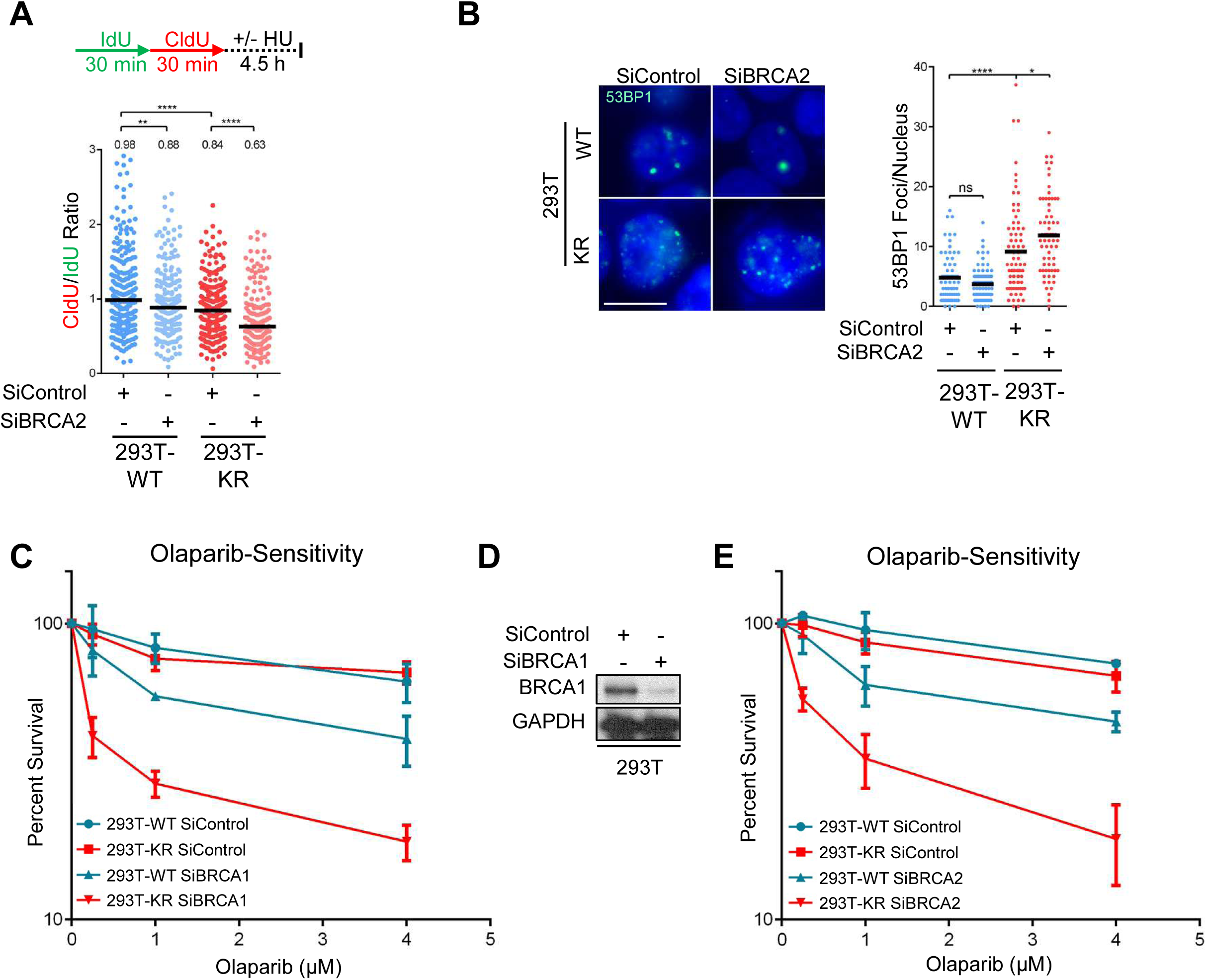
Synergistic interaction between PCNA ubiquitination and the BRCA pathway. **A.** DNA fiber combing assay showing that concomitant loss of BRCA2 and PCNA ubiquitination results in a synergistic increase in nascent strand degradation upon HU-induced fork arrest. The ratio of CldU to IdU tract lengths is presented, with the median values marked on the graph and listed at the top. Asterisks indicate statistical significance. A schematic representation of the fiber combing assay conditions is also presented. **B**. Immunofluorescence experiment showing synergistic increase in 53BP1 foci formation in unsynchronized K164R cells upon BRCA2 depletion. At least 65 cells were quantified for each condition. The mean value is represented on the graphs, and asterisks indicate statistical significance. Representative micrographs are also shown. **C-E**. Clonogenic survival experiments showing that loss of PCNA ubiquitination does not result in olaparib sensitivity, but it drastically increases the olaparib sensitivity of BRCA1 (**C**)- and BRCA2 (**E**)-depleted cells. The average of three experiments, with standard deviations indicated as error bars, is shown. Asterisks indicate statistical significance. Confirmation of BRCA1 knockdown upon siRNA treatment is also shown (**D**).

## Discussion

### PCNA ubiquitination and the regulation of lagging strand replication

Our work uncovered a novel mechanism of replication fork protection controlled by PCNA ubiquitination (Figure S6). We show that PCNA ubiquitination promotes gap filling during normal S-phase. In the absence of PCNA ubiquitination, OF ligation is perturbed, likely because of accumulated gaps on the lagging strand. OF ligation, in turn, permits unloading of PCNA by ATAD5, which enables accurate chromatin assembly by CAF-1. Importantly, we show that defective OF ligation, and the subsequent chromatin assembly deficiency, render cells sensitive to DNA2-mediated nascent tract degradation upon fork stalling and reversal, resulting in double strand break formation and genomic instability. Previously, PCNA SUMOylation has been implicated in regulating chromatin structure at transcribed loci (Li et al., 2018). Our work suggests that this does not affect fork protection, as we show that loss of PCNA ubiquitination, but not of SUMOylation, results in nascent strand degradation.

Our previous work using synthetic genetic array analyses in yeast showed a striking genetic similarity between the PCNA-K164R mutation and inactivation of FEN1 (Rad27 in yeast) and other lagging strand synthesis factors, implicating PCNA ubiquitination in OF maturation (Becker et al., 2015). Moreover, we showed that PCNA ubiquitination was increased in *RAD27* and *CDC9* mutants, and identified a synthetic lethal interaction between the PCNA-K164R mutation and *RAD27* inactivation (Becker et al., 2018; Das-Bradoo et al., 2010; Nguyen et al., 2013) –further indicating that PCNA ubiquitination is required for OFM processing. We show here that human PCNA-K164R cells exhibit hallmarks of lagging strand synthesis defects, and accumulate ssDNA which may directly interfere with OF ligation by LIG1. In yeast, PCNA was shown to be preferentially enriched on the lagging strand during normal DNA replication (Yu et al., 2014). It is thus likely that the single stranded gaps observed in KR cells are on the lagging strand. Indeed, as the lagging strand experiences frequent repriming due to the discontinuous mode of DNA replication, Polδ arrest at endogenous sites of replication stress would result in accumulation of gaps behind the fork as a new DNA synthesis reaction is initiated upon regular repriming of the subsequent OF (Fu et al., 2011; Taylor and Yeeles, 2018). In contrast, stalling of Polε on the leading strand requires a dedicated repriming event which needs to be quickly put in place to resume replication, thus accumulation of gaps on this strand is less likely. We noticed that loss of TLS polymerase Polκ partially phenocopies the fork stability defect in KR cells, suggesting that Polκ is one of the DNA polymerases involved in PCNA ubiquitination-mediated gap filing. However, DNA2 inhibition did not completely rescue the fork protection defect of Polκ-depleted cells, and indeed it was previously shown that loss of this polymerase results in MRE11-mediated fork degradation (Tonzi et al., 2018). This indicates that Polκ also plays a role in fork protection independently of PCNA ubiquitination. In addition, an earlier report proposed that in fission yeast, PCNA ubiquitination enhances interaction with Polδ (Daigaku et al., 2017), suggesting that the activity of the lagging strand replicative polymerase itself may be defective in KR cells.

Interestingly, under normal growth conditions, PCNA-K164R cells show increased fork speed and reduced origin firing. The suppression of origin firing has been shown to result in faster moving forks (Zhong et al., 2013). Interestingly, 53BP1 nuclear bodies were shown to suppress origin firing (Spies et al., 2019). As KR cells show increased 53BP1 foci, we propose that under-replicated DNA in KR cells eventually results in DNA damage and 53BP1 nuclear body accumulation, which in turn suppress origin firing causing the increase in fork speed observed in these cells. A similar mechanism is likely at play in OFM-deficient cells, for example upon knockdown of LIG1 or PARP1, which also show increased fork speed. Interestingly, increased replication fork speed was not reported in K164R mouse embryonic fibroblasts (Vujanovic et al., 2017) suggesting a different control of origin usage in primary cells.

### PCNA ubiquitination and replication fork protection

We show here that inactivation of the ^Ubi^PCNA–LIG1–ATAD5–CAF-1 pathway results in DNA2-mediated nascent strand degradation upon fork reversal. DNA2 was previously shown to degrade stalled forks upon prolonged replication stress in wildtype cells (Thangavel et al., 2015), but in KR cells degradation occurs upon much shorter HU exposure, which does not affect fork stability in wildtype cells. This indicates that PCNA ubiquitination specifically suppresses DNA2-mediated degradation.

Reversal of stalled replication forks is an essential fork stability mechanism. At the same time, reversal renders forks susceptible to MRE11-mediated degradation, unless reversed forks are protected by loading of RAD51 by the BRCA pathway (Ray Chaudhuri et al., 2016; Schlacher et al., 2011; Schlacher et al., 2012; Taglialatela et al., 2017). Depletion of any of the three translocases ZRANB3, HLTF and SMARCAL1 completely restored fork protection in BRCA-deficient cells, indicating that in this genetic context, they work in concert to perform fork reversal (Poole and Cortez, 2017; Taglialatela et al., 2017). In contrast, in KR cells the three translocases have differential impacts. HLTF depletion did not have any effect on fork degradation in KR cells. Besides its translocase activity, HLTF contains a RING ubiquitin ligase domain which catalyzes K63-linked poly-ubiquitination of PCNA at K164, building upon the single ubiquitin moiety initially added by RAD18 (Motegi et al., 2008; Unk et al., 2008). Our findings suggest that HLTF needs to ubiquitinate PCNA in order to perform its translocase activity. ZRANB3 had a moderate impact on fork protection. Although ZRANB3 preferentially binds poly-ubiquitinated PCNA, it is able to interact with unmodified PCNA through its PIP-box (Ciccia et al., 2012), thereby explaining its intermediate phenotype. Lastly, depletion of SMARCAL1 completely suppressed fork degradation, indicating that SMARCAL1 is the primary fork reversal activity operating in KR cells. Our findings demonstrate that the three translocases do not necessarily have to act together in fork protection, but instead the particular fork composition dictates which of them performs the fork reversal process. While nascent strand degradation described in BRCA-deficient cells involves the activity of MRE11, we show here that resection of stalled forks in KR cells is performed by DNA2. Not only are the nucleases different, but the primary defects that enable degradation are distinct. In PCNA-K164R cells, RAD51 loading on reversed forks is intact. Instead, we propose that the aberrant nucleosome deposition promotes nascent tract degradation by DNA2. DNA2 is also involved in fork degradation in BRCA-deficient cells (Ray Chaudhuri et al., 2016), but that activity is performed in conjunction with MRE11 and thus is different than what we report here in PCNA-K164R cells, where we find no evidence of MRE11 activity.

In order for nascent tract degradation to be detectable by the DNA fiber combing assay, both nascent strands in symmetrical reversed forks must be degraded. It was previously shown that, during DSB resection, in order to generate the 3’ overhangs, MRE11 initiates a nick on the complementary strand, followed by resection in a 3’-5’ manner towards the DSB, thereby allowing access to long-range 5’-3’ nucleases (Garcia et al., 2011; Paull and Gellert, 1998). It is thus conceivable that that in BRCA-deficient cells that lack RAD51-mediated protection of reversed forks, MRE11 is able to attack both nascent strands using its 3’-5’ nuclease activity. In contrast, DNA2 can only perform resection in the 5’-3’ direction. Asymmetrical reversed forks presenting with a 5’ss-DNA overhang may represent a substrate for DNA2, which would yield quantifiable degradation in the DNA fiber combing assay. Indeed, aside from DSB resection, DNA2 is capable of processing long 5’ss-DNA flaps arising from excessive strand displacement during OF synthesis (Ayyagari et al., 2003; Fortini et al., 2011; Rossi et al., 2018). It is possible that reversed forks with 5’-overhangs arising from the lagging strand may resemble 5’ss-DNA flaps observed during OFM.

How might the asymmetry between leading and lagging strands in PCNA-K164R arise? OFM defects in these cells result in retention of PCNA on chromatin, which has been proposed to sequester CAF-1 resulting in aberrant chromatin assembly (Janke et al., 2018). In line with this, loss of PCNA ubiquitination shows a similar impact on nucleosomal compaction as CHAF1A depletion, suggesting that CAF-1 sequestration is functionally equivalent with loss of its activity. Nucleosome positioning was shown to have a direct impact on OF periodicity and disrupting chromatin assembly by inactivating CAF-1 results in abnormally long OFs (Smith and Whitehouse, 2012; Yadav and Whitehouse, 2016). Moreover, nucleosomal positioning guides the priming activity of the Polα-primase (Kurat et al., 2017). Therefore, we speculate that in KR cells, replication forks encounter a sparse chromatin organization due to impaired CAF-1 activity, which alters the periodicity of OF priming. This would give rise to longer OFs that initiate further ahead of leading strand synthesis, resulting in 5’-overhangs when fork reversal occurs (Figure S6).

### PCNA ubiquitination and the response to PARP inhibitors

PARP inhibitors such as olaparib are effective in the clinical treatment of BRCA-mutant breast and ovarian carcinomas (Moore et al., 2018). However, tumors eventually acquire resistance to these drugs. Replication fork protection is considered an important component of PARPi resistance in BRCA-deficient cells (Ray Chaudhuri et al., 2016; Taglialatela et al., 2017). We show here that concomitant loss of PCNA ubiquitination and the BRCA pathway results in synergistic increase in fork degradation, DNA damage accumulation, and olaparib sensitivity. These findings suggest that PCNA ubiquitination provides a survival mechanism for BRCA-deficient cells exposed to DNA damaging agents, and its inactivation may potentiate the anti-tumor effect of PARPi in these cells.

Moreover, our work sheds new light on the mechanism involved in the synthetic lethality between BRCA-deficiency and PARPi (Bryant et al., 2005; Farmer et al., 2005). Recent studies revealed a previously unknown role of PARP1 as a sensor of unligated OFs during unperturbed S-phase (Hanzlikova et al., 2018). Therefore, it is possible that a major effect of PARPi is preventing the ligation of OFs that have failed conventional ligation by LIG1. Yet it has remained unclear whether perturbed OF ligation causes toxicity in BRCA-deficient cells. In this study we show that loss of PCNA ubiquitination, which causes defects in OF ligation due to inefficient gap-filling behind progressing forks, synergistically exacerbates the sensitivity of BRCA-deficient cells to PARPi. This indicates that cells with intact BRCA1/2 function are able to tolerate unligated OFs, as opposed to BRCA-mutant cells where the perturbation of OF ligation through PARPi and the concomitant loss of PCNA ubiquitination is increasingly toxic. These observations point towards perturbation of OF ligation as a potential mechanism underlying the synthetic lethality of PARPi in cells with BRCA deficiency and reflect the importance of PCNA-ubiquitination in mediating tolerance to PARPi in these cells.

## Acknowledgments

We would like to thank Drs. Alessandro Vindigni, Alberto Ciccia, James Broach and Mark Hedglin for materials and advice; and the Penn State College of Medicine Flow Cytometry and Imaging cores. This work was supported by: NIH R01GM134681 (to AKB and GLM); NIH R01ES026184 (to GLM); NIH 5R01GM074917 (to AKB); NIH R01CA085344 (to BS).

## Methods

### Cell culture and protein techniques

Human 293T, RPE1 and U2OS cells were grown in DMEM supplemented with 10% Fetal Calf Serum.

To generate the K164R cells, the gRNA sequences used were: TTTCACTCCGTCTTTTGCACAGG for 293T cells and GCAAGTGGAGAACTTGGAAATGG for RPE1 cells. The sequences were cloned into the pX458 vector (pSpCas9BB-2A-GFP; obtained from Addgene). Cells were co-transfected with this vector and a repair template spanning the K164 genomic locus but containing the K164R mutation (AAA-AGA codon change). Transfected cells were FACS-sorted into 96-well plates using a BD FACSAria II instrument. Resulting monoclonal cultures were screened by Western blot for loss of PCNA ubiquitination using an antibody specific for this modification. For verification of positive cell lines, the targeted genomic region was PCR amplified from genomic DNA, cloned into pBluescript, and multiple clones were Sanger-sequenced to ensure that all alleles are identified. For exogenous PCNA expression, pLV-puro-CMV lentiviral constructs encoding wildtype or the K164R variant were obtained from Cyagen. Infected cells were selected by puromycin.

For RAD18 gene knockout, the commercially available CRISPR/Cas9 KO Plasmid BRCA2 CRISPR/Cas9 KO plasmid was used (Santa Cruz Biotechnology sc-406099). Transfected cells were FACS-sorted into 96-well plates using a BD FACSAria II instrument. Resulting colonies were screened by Western blot.

Denatured whole cell extracts were prepared by boiling cells in 100 mM Tris, 4% SDS, 0.5M β-mercaptoethanol. Chromatin fractionation was performed as previously described (Wysocka et al., 2001). For poly(ADP-ribose) glycohydrolase (PARG) inhibition, cells were incubated with 10μM PDD00017273 (Sigma SML1781) for 45min prior to harvesting, as previously described for detection of OFM defects (Hanzlikova et al., 2018).

Antibodies used for Western blot were:

PCNA (Cell Signaling Technology 2586)

Ubiquityl-PCNA Lys164 (Cell Signaling Technology 13439)

Chk2 (Cell Signaling Technology 2662)

pChk2-T68 (Cell Signaling Technology 2661)

Chk1 (Cell Signaling Technology 2360)

pChk1-317 (Cell Signaling Technology 2344)

BRCA2 (Bethyl A303-434A)

EXO1 (Santa Cruz Biotechnology sc-56092)

WRN (Santa Cruz Biotechnology sc-5926)

CTIP (Santa Cruz Biotechnology sc-271339)

MUS81 (Santa Cruz Biotechnology sc-47692)

RAD18 (Cell Signaling Technology 9040)

UBC9 (Santa Cruz Biotechnology sc-10759)

RAD51 (Santa Cruz Biotechnology sc-8349)

ZRANB3 (Bethyl A303-033A)

HLTF (Santa Cruz Biotechnology sc-398357)

SMARCAL1 (Santa Cruz Biotechnology sc-376377)

RECQL1 (Santa Cruz Biotechnology sc-166388)

PAR chains (ENZO ALX-804-220)

LaminB1 (Abcam ab16048)

LIG1 (Santa Cruz Biotechnology sc-271678)

PARP1 (Cell Signaling Technology 8542)

REV1 (Santa Cruz Biotechnology sc-393022)

POLK (Santa Cruz Biotechnology sc-166667)

CHAF1A (Cell Signaling Technology 5480)

BRCA1 (Santa Cruz Biotechnology sc-642)

Vinculin (Santa Cruz Biotechnology sc-73614)

GAPDH (Santa Cruz Biotechnology sc-47724)

For gene knockdown, cells were transfected with Stealth siRNA (Life Tech) using Lipofectamine RNAiMAX reagent. The siRNA targeting sequences used were:

BRCA2: GAGAGGCCTGTAAAGACCTTGAATT

EXO1: CCTGTTGAGTCAGTATTCTCTTTCA

WRN: TGGGCTCCTGCAGACATTAACTTAA

CTIP: GGGTCTGAAGTGAACAAGATCATTA

MUS81: TTTGCTGGGTCTCTAGGATTGGTCT

RAD18: CATATTAGATGAACTGGTATT

UBC9: TAAACAAGCCTCCTTCCCACGGAGT

RAD51: CCATACTGTGGAGGCTGTTGCCTAT

HLTF: TGCATGTGCATTAACTTCATCTGTT

ZRANB3: TGGCAATGTAGTCTCTGCACCTATA

SMARCAL1: CACCCTTTGCTAACCCAACTCATAA

RECQL1: ACAGGAGGUGGAAAGAGCTTATGTT

LIG1: CCAAGAACAACTATCATCCCGTGGA

PARP1: AAACATGGGCGACTGCACCATGATG

REV1: GAAATCCTTGCAGAGACCAAACTTA

POLK: CAGCCATGCCAGGATTTATTGCTAA

ATAD5: GGTACGCTTTAAGACAGTTACTGTT

CHAF1A: GCCTGAATCTTGTCCCAAATT

BRCA1: AATGAGTCCAGTTTCGTTGCCTCTG

### Functional assays

For clonogenic experiments, 1000 cells were seeded in 6-well plates. For UV sensitivity, cells were treated 24 h after seeding. For cisplatin and olaparib treatment, cells were seeded in indicated drug concentrations for 24 and 72 h respectively, followed by media change. Two weeks later, colonies were stained with Crystal violet.

For time-course proliferation experiments, 500 cells were seeded in wells of 96-well plates, and cellular viability was scored at indicated days using the CellTiterGlo reagent (Promega G7572).

EdU incorporation was assayed using the Click-iT Plus kit (Invitrogen C10633) according to the manufacturer’s instructions.

Translesion synthesis SupF assay was previously described (Wang et al., 1995). Cells were transfected with UVC-irradiated (1000J/m2) pSP189 (SupF) plasmid. Three days later, the plasmid was recovered using a miniprep kit (Promega), DpnI digested and transformed into MBM7070 indicator bacteria. Transformants were selected on plates containing 1mM IPTG and 100μg/ml X-gal. The ratio of white (mutant) to total (blue + white) colonies was scored as mutation frequency.

### BrdU alkaline comet assay

Cells were incubated with 100μM BrdU for 15min, followed by media removal, PBS wash, and incubation in fresh media for 1h. Cells were harvested and subjected to the alkaline comet assay using the CometAssay kit (Trevigen 4250-050) according to the manufacturer instructions. Slides were stained with primary anti-BrdU (BD 347580) and secondary Alexa Fluor 568 (Invitrogen A11031) antibodies. Slides were mounted with DAPI-containing Vectashield mounting medium (Vector Labs) and imaged using a Leica SP5 confocal microscope. The percent tail DNA was calculated using CometScore 2.0 software.

### DNA Fiber Assay

Cells were incubated consecutively with 100µM CldU and 100µM IdU for the indicated times. Hydroxyurea and nuclease inhibitors (50μM Mirin for MRE11 inhibition or 30μM C5 (Liu et al., 2016) for DNA2 inhibition) were added as indicated. Next, cells were harvested and DNA fibers were obtained using the FiberPrep kit (Genomic Vision). DNA fibers were stretched on glass coverslips using the FiberComb Molecular Combing instrument (Genomic Vision). Slides were incubated with primary antibodies (Abcam 6326 for detecting CIdU; BD 347580 for detecting IdU; Millipore Sigma MAB3034 for detecting DNA), washed with PBS, and incubated with Cy3, Cy5, or BV480 –coupled secondary antibodies (Abcam 6946, Abcam 6565, BD Biosciences 564879). Following mounting, slides were imaged using a Leica SP5 confocal microscope. At least 100 tracts were quantified for each sample.

### Immunofluorescence

Cells were seeded on sterile glass coverslips coated with Poly-L-Lysine (Sigma P8920) as per manufacturer’s instructions and allowed to incubate for 24h. For RPA and 53BP1 foci detection, cells were fixed with 3.7% paraformaldehyde for 15min, followed by three washes with PBS. Cells were then permeabilized with 0.5% Triton X-100 for 10min. After two washes with PBS, slides were blocked with 3% BSA in PBS for 15min, followed by incubation with the primary antibody diluted in 3% BSA in PBS, for 2h at room temperature. After three washes with PBS, the secondary antibody was added for 1h. Slides were mounted with DAPI-containing Vectashield mounting medium (Vector Labs). For PCNA foci detection, cells were pre-extracted with 0.5% Triton X-100, followed by one PBS wash and methanol fixation for 30min at −20ᵒC. After two washes with PBS, cells were blocked with 3% BSA in PBS for 15min. Primary and secondary antibody treatments as well as mounting were performed as mentioned above. Primary antibodies used for immunofluorescence were: 53BP1 (Bethyl A300-272A); RPA32 (Abcam ab2175); PCNA (Cell Signaling Technology 2586). Secondary antibodies used were AlexaFluor 488 or AlexaFluor 568 (Invitrogen A11001, A11008, A11031, and A11036). Slides were imaged using a Leica SP5 confocal microscope. The number of foci/nucleus was quantified using ImageJ software. For the PCNA immunofluorescence, in order to remove non-S-phase cells, only cells with at least 2 foci were quantified, corresponding to the top three quartiles in wildtype, were included in the quantification.

### Quantification of gene expression by real-time quantitative PCR (RT-qPCR)

Total mRNA was purified using TRIzol reagent (Life Tech). To generate cDNA, 1μg RNA was subjected to reverse transcription using the RevertAid Reverse Transcriptase Kit (Thermo Fisher Scientific) with oligo-dT primers. Real-time qPCR was performed with PerfeCTa SYBR Green SuperMix (Quanta), using a CFX Connect Real-Time Cycler (BioRad). The cDNA of GAPDH gene was used for normalization. Primers used were:

RADX for: ATGATGTGACGATCTCAGATGGG

RADX rev: CCCCTGGCCTATCCTTTTCTC

ATAD5 for: AGGAAGAGATCCAACCAACG

ATAD5 rev: ATGTTTCGAAGGGTTGGCAG

GAPDH for: AAATCAAGTGGGGCGATGCTG;

GAPDH rev: GCAGAGATGATGACCCTTTTG

### Micrococcal nuclease assay

Cells were trypsinized, washed with PBS and lysed with cold NP-40 lysis buffer (10mM Tris-HCL, 10mM NaCl, 3mM MgCl2, 0.5% NP-40, 0.15mM spermine and 0.5mM spermidine) for 5min. The resulting nuclei were washed once and resuspended in MNase digestion buffer (10mM Tris-HCl, 15mM NaCl, 60mM KCl, 1mM CaCl_2_, 0.15mM spermine and 0.5mM spermidine). Resuspended cells were digested with 2.5 U MNase in digestion buffer for the indicated times. The reaction was stopped by adding an equal volume of the MNase stop buffer (100mM EDTA, 10mM EGTA, 15mM NaCl, 60mM KCl and 2% SDS) followed by proteinase K digestion (final concentration of 0.0375μg/μL) at 37ᵒC overnight. DNA was isolated using phenol-chloroform extraction and subsequently subjected to RNAse A digestion. Samples were run on 1.2% agarose gels and visualized using Gel Red. Signal intensities were quantified using Icy software (Institut Pasteur).

### Statistical analyses

For the DNA fiber and immunofluorescence experiments, the Mann-Whitney statistical test was performed. For all other assays, the statistical analysis performed was the TTEST (two-tailed, unequal variance). Statistical significance is indicated for each graph (ns = not significant, for P > 0.05; * for P < 0.05; ** for P < 0.01; *** for P < 0.001; **** for P < 0.0001).

## Legends to Supplemental Figures

**Figure S1.**
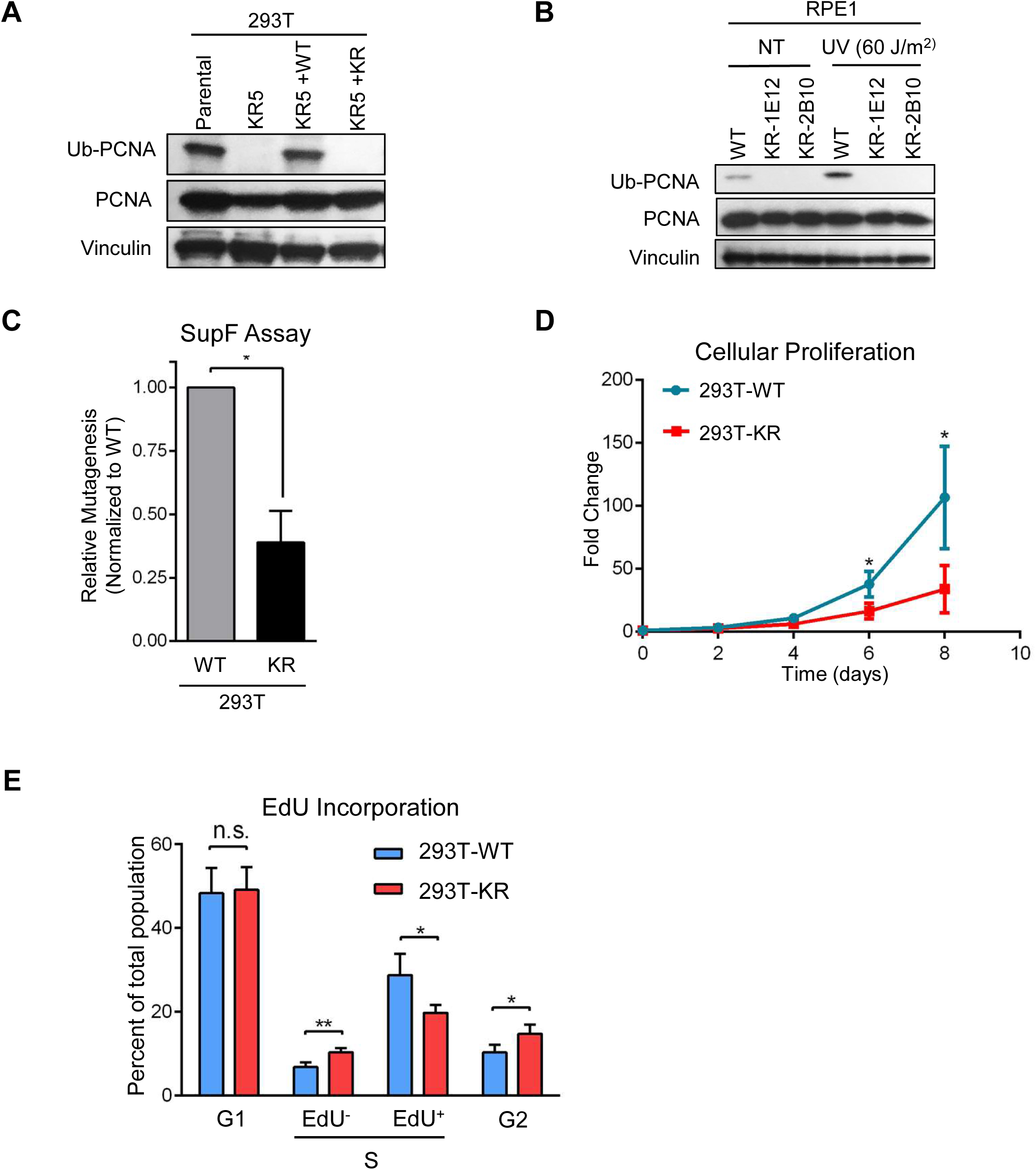
Functional characterization of human PCNA-K164R mutant cells. **A**. Western blot showing PCNA and ubiquitinated PCNA levels in parental and the KR5 clone obtained from CRISPR/Cas9-mediated genetic editing of the PCNA alleles in 293T cells. Denatured whole cell extracts are shown. The KR5 clone was corrected be re-expression of wildtype or K164R PCNA from a lentiviral construct, to restore normal PCNA levels. **B**. Western blot showing the loss of PCNA ubiquitination in two different RPE1-K164R clones generated through CRISP/Cas9 technology. Denatured whole cell extracts of cells under normal growth conditions, or 3h after exposure to the indicated UV dose, were analyzed. **C**. SupF shuttle plasmid mutagenesis assay showing reduced UV-induced mutagenesis in 293T-K164R cells. The average of three experiments, with standard deviations indicated as error bars, is shown. Asterisks indicate statistical significance. **D**. Cellular proliferation experiment showing reduced growth rate of 293T-K164R cells under normal growth conditions. The average of three experiments, with standard deviations indicated as error bars, is shown. Asterisks indicate statistical significance. **E**. EdU incorporation assay showing reduced proportion of cells actively undergoing DNA synthesis under normal growth conditions. The average of three experiments, with standard deviations indicated as error bars, is shown. Asterisks indicate statistical significance.

**Figure S2.**
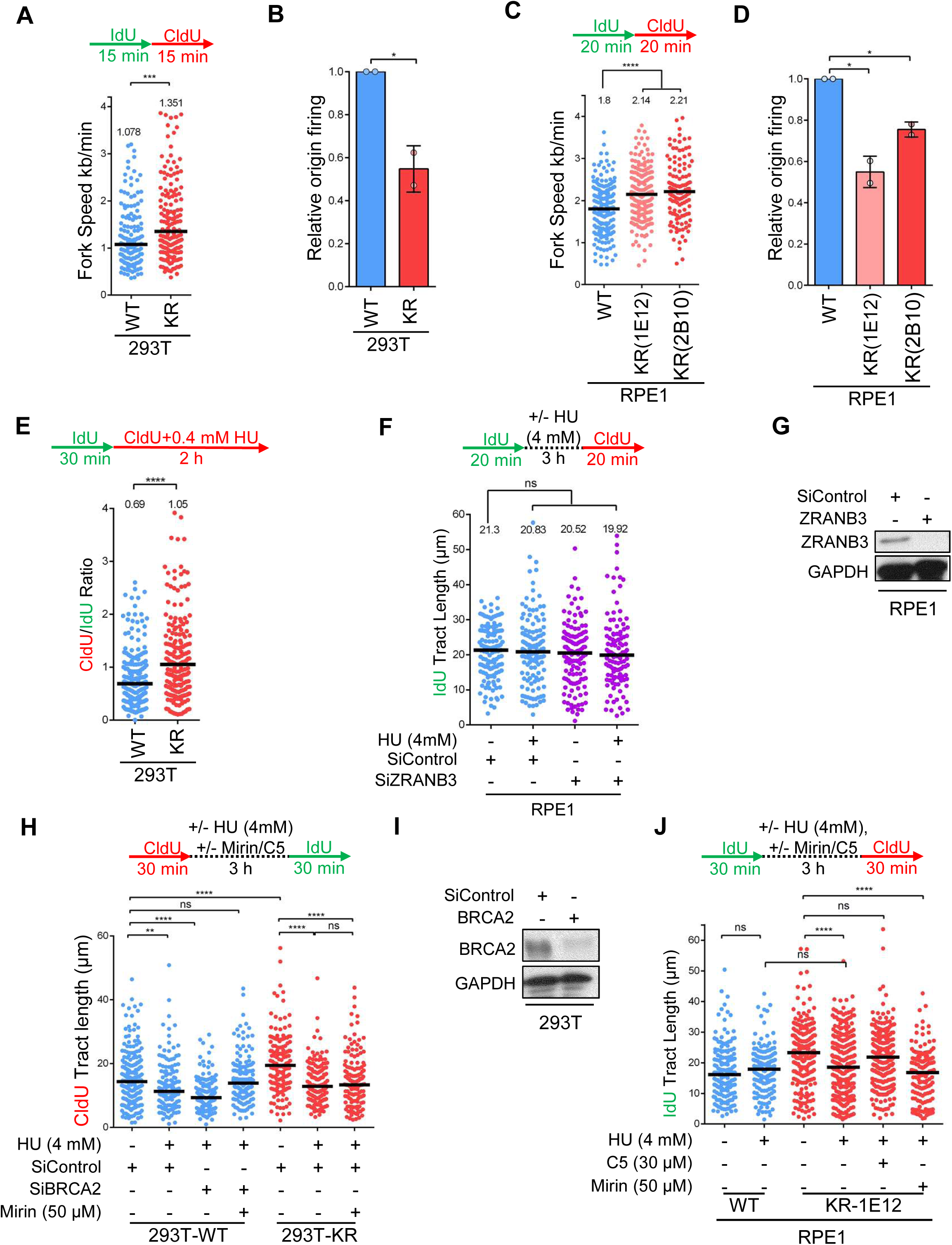
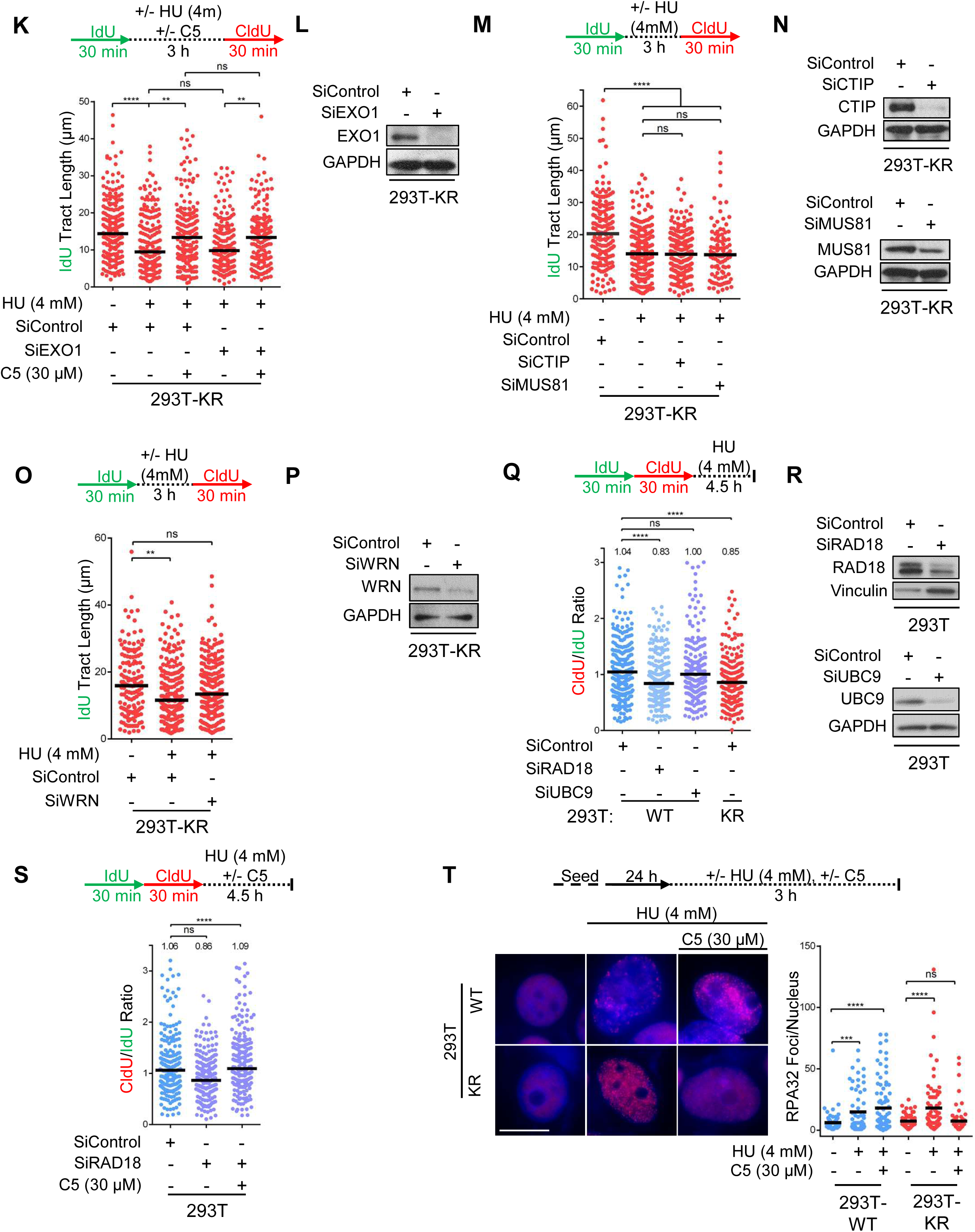
Loss of PCNA ubiquitination results in faster replication forks, which are susceptible to DNA2-mediated degradation upon replication stress. **A-D**. DNA fiber combing experiments showing increased fork speed and reduced origin firing in 293T-K164R (**A, B**) and RPE1-K164R (**C, D**) cells. For fork speed measurements (**A, C**), each 1μm in fiber length was set to correspond to 2 kilobases of DNA. Median values are marked on the graphs and listed at the top. Asterisks indicate statistical significance. Schematic representations of the assay conditions are also presented. Origin firing (**B, D**) was determined by calculating the ratio of the number of CldU-only tracts to that of tracts with adjacent IdU and CldU labeling. The average of two experiments, with standard deviations indicated as error bars, is shown. The individual values are also presented. Asterisks indicate statistical significance. **E**. DNA fiber combing experiment showing reduced fork slowing in 293T-K164R cells upon low-level replication stress exposure. The ratio of CldU to IdU tract lengths is presented, with the median values marked on the graph and listed at the top. Asterisks indicate statistical significance. A schematic representation of the assay conditions is also presented. **F**. ZRANB3 depletion does not affect fork speed under normal growth conditions, or nascent strand degradation upon fork arrest. The quantification of the IdU tract length is shown, with the median values marked on the graph and listed at the top. Statistical significance, and a schematic representation of the DNA fiber combing assay conditions, are also presented. **G**. Western blot showing ZRANB3 depletion upon siRNA-mediated knockdown. **H**. MRE11 inhibition by mirin suppresses HU-induced nascent strand degradation in BRCA2-knockdown cells, but not in 293T-K164R cells. The quantification of the CldU tract length is shown, with the median values marked on the graph. Statistical significance, and a schematic representation of the DNA fiber combing assay conditions, are also presented. **I**. Western blot showing BRCA2 depletion upon siRNA-mediated knockdown. **J**. DNA2 inhibition by C5, but not MRE11 inhibition by mirin, suppresses HU-induced nascent strand degradation in RPE1-K164R cells. The quantification of the IdU tract length is shown, with the median values marked on the graph. Statistical significance, and a schematic representation of the DNA fiber combing assay conditions, are also presented. **K**. DNA2 inhibition by C5, but not EXO1 depletion, suppresses HU-induced nascent strand degradation in 293T-K164R cells. The quantification of the IdU tract length is shown, with the median values marked on the graph. Statistical significance, and a schematic representation of the DNA fiber combing assay conditions, are also presented. **L**. Western blot showing EXO1 depletion upon siRNA-mediated knockdown. **M**. Loss of CTIP or MUS81 does not suppress the HU-induced nascent strand degradation in 293T-K164R cells. The quantification of the IdU tract length is shown, with the median values marked on the graph. Statistical significance, and a schematic representation of the DNA fiber combing assay conditions, are also presented. **N**. Western blot showing CTIP and MUS81 depletion upon siRNA-mediated knockdown. **O**. Depletion of WRN partially suppresses the HU-induced nascent strand degradation in 293T-K164R cells. The quantification of the IdU tract length is shown, with the median values marked on the graph. Statistical significance, and a schematic representation of the DNA fiber combing assay conditions, are also presented. **P**. Western blot showing WRN depletion upon siRNA-mediated knockdown. **Q.** Depletion of RAD18, but not of UBC9, results in HU-induced nascent strand degradation. The ratio of CldU to IdU tract lengths is presented, with the median values marked on the graph and listed at the top. Asterisks indicate statistical significance. A schematic representation of the DNA fiber combing assay conditions is also presented. **R.** Western blot showing RAD18 and UBC9 depletion upon siRNA-mediated knockdown. **S.** Depletion of RAD18 in wildtype 293T cells results in HU-induced nascent strand degradation mediated by DNA2. The ratio of CldU to IdU tract lengths is presented, with the median values marked on the graph and listed at the top. Asterisks indicate statistical significance. A schematic representation of the DNA fiber combing assay conditions is also presented. **T**. Immunofluorescence experiment showing that HU induced RPA32 foci formation is suppressed by DNA2 inhibition in 293T-K164R cells, but not in wildtype cells. At least 75 cells were quantified for each condition. The mean value is represented on the graphs, and asterisks indicate statistical significance. Representative micrographs are also shown.

**Figure S3.**
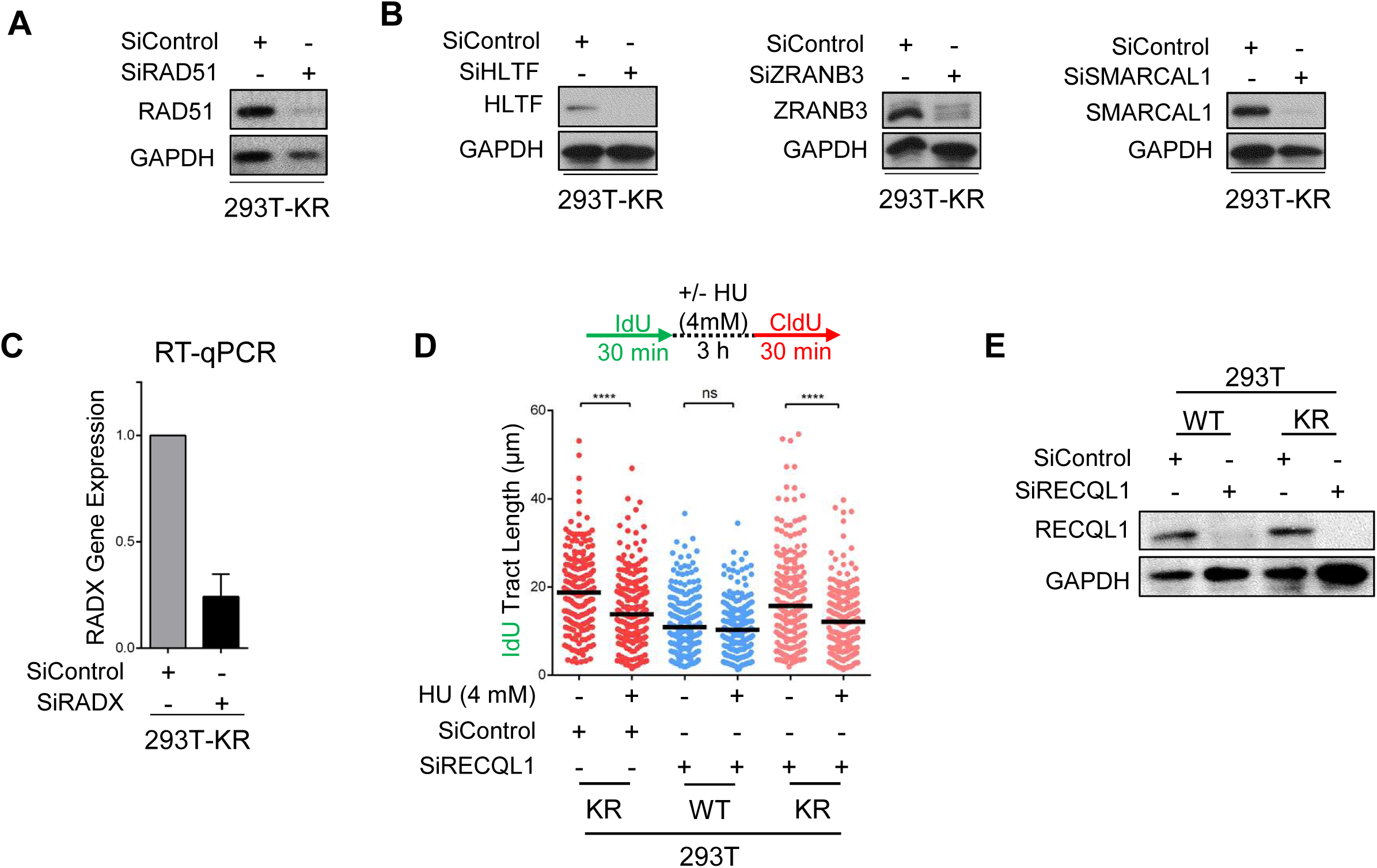
Impact of fork reversal and restart on HU-induced nascent tract degradation in PCNA-K164R cells. **A**. Western blot confirming RAD51 depletion upon siRNA-mediated knockdown. **B.** Western blots showing depletion of HLTF, ZRANB3 and SMARCAL1 upon siRNA-mediated knockdown. **C**. RT-qPCR experiment showing reduction in RADX mRNA levels upon siRNA-mediated knockdown. The average of two technical replicates is shown, with error bars indicating standard deviations. (No antibody was available to us for verifying the depletion by Western blot). **D.** RECQL1 is not involved in the nascent tract degradation observed in K164R cells upon HU exposure, as its knockdown does not induce fork degradation in wildtype cells, and does not the affect the degradation observed in KR cells. The quantification of the IdU tract length is presented, with the median values marked on the graph. Asterisks indicate statistical significance. A schematic representation of the assay conditions is also presented. **E**. Western blot confirming RECQL1 depletion upon siRNA-mediated knockdown.

**Figure S4.**
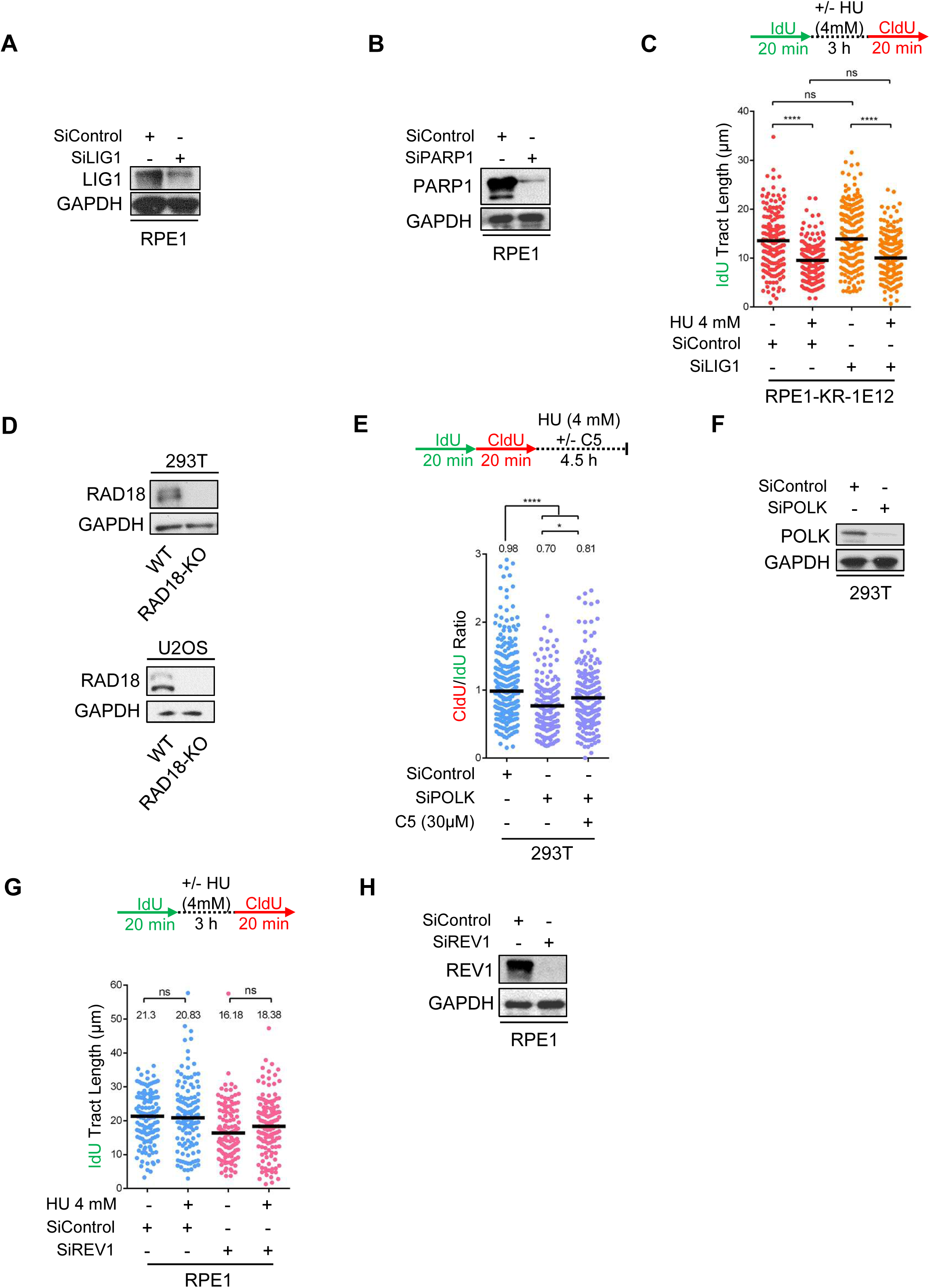
Impact of Okazaki fragment maturation factors and TLS polymerases on replication fork protection. **A**. Western blot confirming LIG1 depletion upon siRNA-mediated knockdown. **B**. Western blot confirming PARP1 depletion upon siRNA-mediated knockdown. **C**. DNA fiber combing assay showing that LIG1 depletion in RPE1 cells does not further increase fork progression rate and HU-induced degradation in K164R cells. The quantification of the IdU tract length is presented, with the median values marked on the graph. Asterisks indicate statistical significance. A schematic representation of the assay conditions is also presented. **D.** Western blots showing the loss of RAD18 expression in 293T and U2OS RAD18-knockout cells. **E.** POLK depletion results in HU-induced nascent strand degradation which is partially dependent on DNA2 enzymatic activity. The ratio of CldU to IdU tract lengths is presented, with the median values marked on the graph and listed at the top. Asterisks indicate statistical significance. A schematic representation of the fiber combing assay conditions is also presented. **F**. Western blot confirming POLK depletion upon siRNA-mediated knockdown. **G**. DNA fiber combing assay showing that REV1 depletion does not cause degradation of arrested replication forks. The quantification of the IdU tract length is presented, with the median values marked on the graph and listed at the top. Asterisks indicate statistical significance. A schematic representation of the assay conditions is also presented. **H**. Western blot confirming REV1 depletion upon siRNA-mediated knockdown.

**Figure S5.**
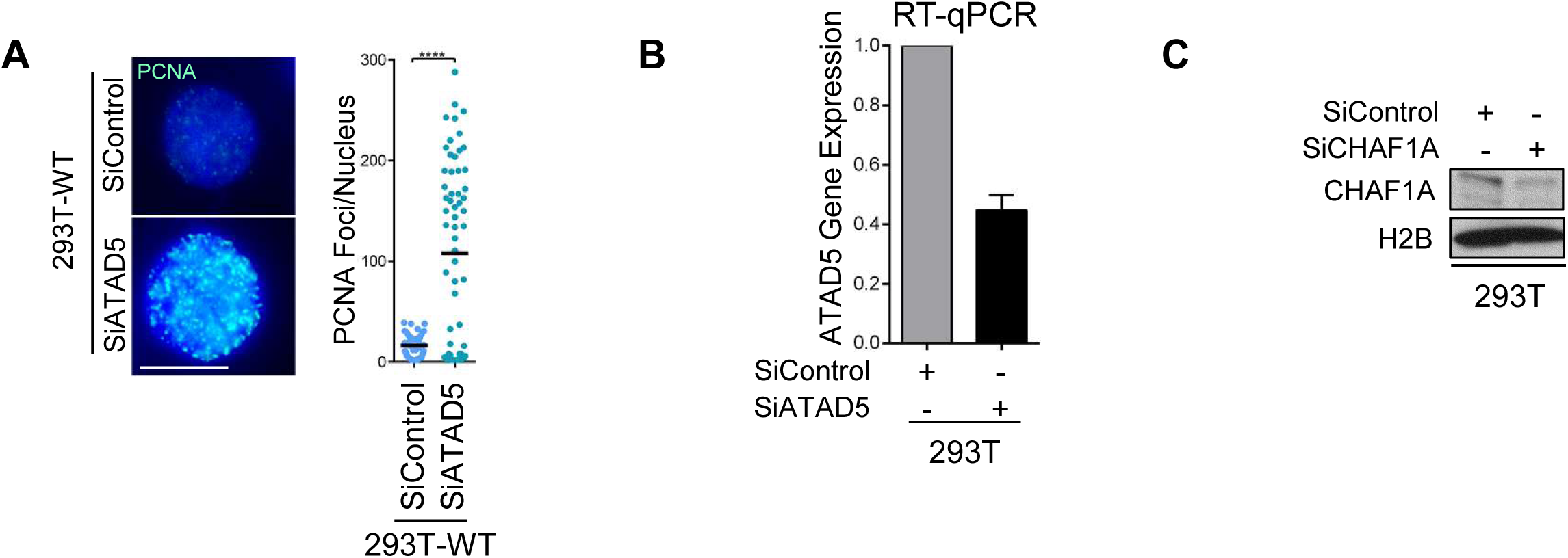
Impact of PCNA chromatin retention and CHAF1A-mediated chromatin assembly on replication fork protection. **A.** PCNA immunofluorescence showing increased PCNA retention on chromatin upon ATAD5 depletion. At least 65 cells were quantified for each condition. The mean value is represented on the graphs, and asterisks indicate statistical significance. Representative micrographs are also shown. **B**. RT-qPCR experiment showing reduction in ATAD5 mRNA levels upon siRNA-mediated knockdown. The average of two technical replicates is shown, with error bars indicating standard deviations. (No antibody was available to us for verifying the depletion by Western blot). **C**. Western blot confirming CHAF1A depletion upon siRNA-mediated knockdown.

**Figure S6.**
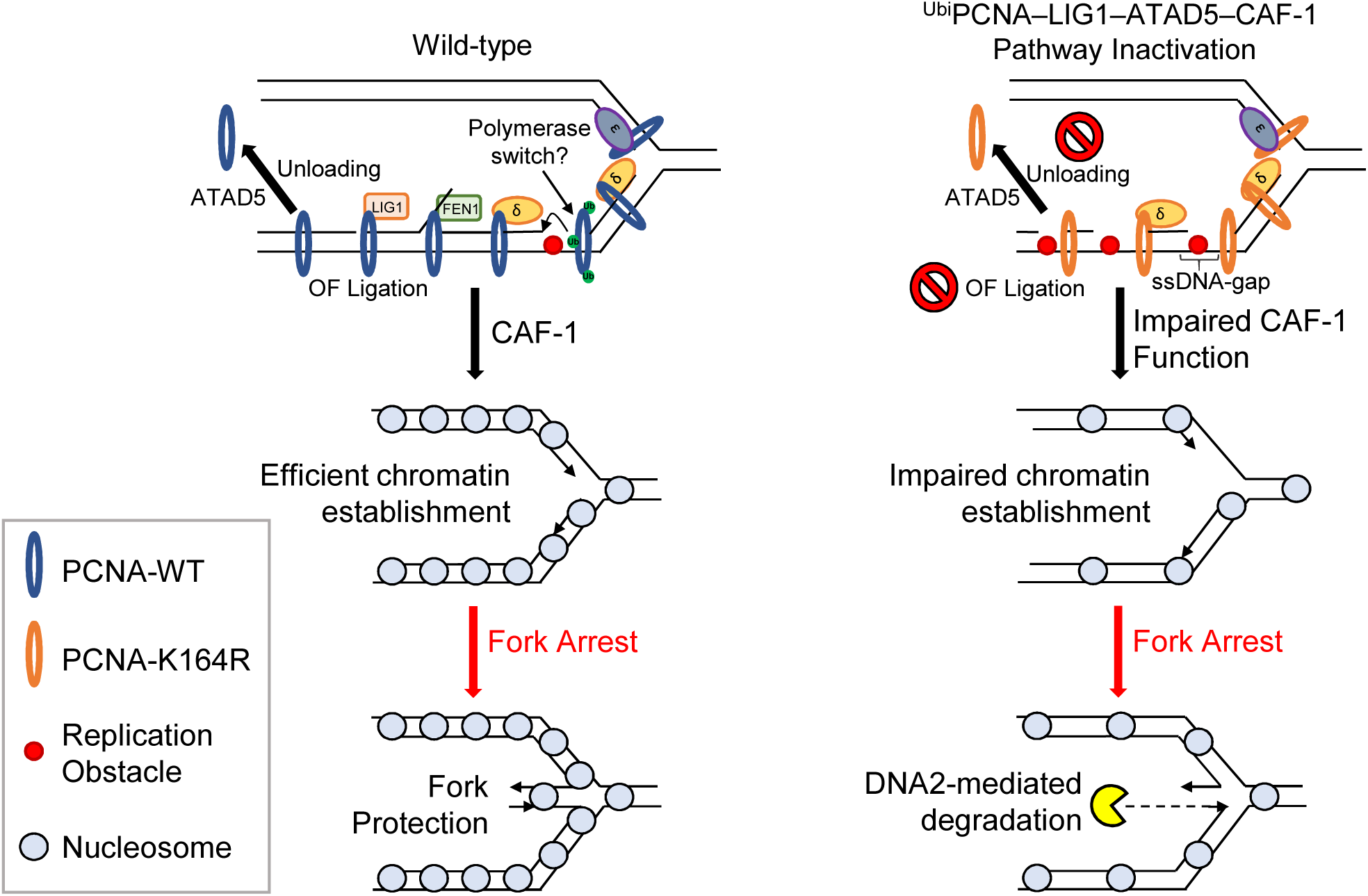
Model depicting the ^Ubi^PCNA–LIG1–ATAD5–CAF-1 pathway. PCNA ubiquitination ensures efficient OF ligation by mediating gap-filling behind progressing replication forks. A deficiency in PCNA-ubiquitination interferes with complete OF synthesis and ligation, thereby precluding ATAD5-mediated PCNA unloading from the lagging strand. This disturbs the efficiency of chromatin establishment by CAF-1 which results in replication forks encountering a sparse chromatin organization. The altered arrangement of nucleosomes directly affects Okazaki fragment priming and length, giving rise to a 5’-overhang in the regressed arms when fork reversal occurs. We propose that this altered reversed fork structure is a preferred substrate for DNA2, leading to uncontrolled resection.

